# A user-friendly tool for cloud-based whole slide image segmentation, with examples from renal histopathology

**DOI:** 10.1101/2021.08.16.456524

**Authors:** Brendon Lutnick, David Manthey, Jan U. Becker, Brandon Ginley, Katharina Moos, Jonathan E. Zuckerman, Luis Rodrigues, Alexander J. Gallan, Laura Barisoni, Charles E. Alpers, Xiaoxin X. Wang, Komuraiah Myakala, Bryce A. Jones, Moshe Levi, Jeffrey B. Kopp, Teruhiko Yoshida, Seung Seok Han, Sanjay Jain, Avi Z. Rosenberg, Kuang Yu. Jen, Pinaki Sarder, the Kidney Precision Medicine Project

## Abstract

**Background:** Image-based machine learning tools hold great promise for clinical applications in nephropathology and kidney research. However, the ideal end-users of these computational tools (e.g., pathologists and biological scientists) often face prohibitive challenges in using these tools to their full potential, including the lack of technical expertise, suboptimal user interface, and limited computation power.

**Methods:** We have developed *Histo-Cloud*, a tool for segmentation of whole slide images (WSIs) that has an easy-to-use graphical user interface. This tool runs a state-of-the-art convolutional neural network (CNN) for segmentation of WSIs in the cloud and allows the extraction of features from segmented regions for further analysis.

**Results:** By segmenting glomeruli, interstitial fibrosis and tubular atrophy, and vascular structures from renal and non-renal WSIs, we demonstrate the scalability, best practices for transfer learning, and effects of dataset variability. Finally, we demonstrate an application for animal model research, analyzing glomerular features in murine models of aging, diabetic nephropathy, and HIV associated nephropathy.

**Conclusion:** The ability to access this tool over the internet will facilitate widespread use by computational non-experts. *Histo-Cloud* is open source and adaptable for segmentation of any histological structure regardless of stain. Histo-Cloud will greatly accelerate and facilitate the generation of datasets for machine learning in the analysis of kidney histology, empowering computationally novice end-users to conduct deep feature analysis of tissue slides.

## INTRODUCTION

Recent advances in machine learning techniques have led to previously unachievable performance for image analysis tasks. In particular, convolutional neural networks (CNNs)^1^, a form of deep learning, have great potential for impactful applications in computational analysis of image structures. Successful adoption of these tools to biomedical image data promises a paradigm shift in both biological science and healthcare^2^.

In the field of pathology, the practice of digitizing histological slides has become common practice^3^, facilitating the application of CNNs for analysis. Digitally scanned histology slides, known as whole slide images (WSIs), are often gigapixels in size. Due to the size of these images and the diversity of structures that can be present, parsing WSIs into biologically relevant sub-compartments is often an important first step for tissue analysis^4^.

CNNs have been successfully utilized by many research groups for the segmentation of WSIs^4-9^. However, thus far tools to segment WSIs have been complex to deploy and use, requiring knowledge of the command line interface and computational expertise^10-12^. The ideal user for these tools is the pathologist or biological scientist, whose clinical workflow or research questions could benefit from fast and accurate segmentation of relevant structures^2^.

To address this gap, we have developed *Histo-Cloud*, a powerful tool for the segmentation of WSIs and deployed it as a suite of easy-to-use plugins using the Digital Slide Archive (DSA)^13^, an open source cloud-based WSI repository with a built in slide viewer. *Histo-Cloud* was designed with flexibility in mind and is agnostic to tissue type or structure. Segmentation of new structures of interest is possible by retraining the CNN used for segmentation, which can be conveniently performed within the cloud interface.

## RESULTS

To demonstrate *Histo-Cloud*’s performance characteristics and segmentation potential, a variety of segmentation tasks from renal biopsy WSIs were tested. For each task, performance was evaluated on holdout WSIs and independent test slides selected from datasets never used for training. A description of the datasets used for the studies below, including sources, disease pathology, tissue thickness, staining, and image acquisition is available in the *Methods* section and is summarized in ***Table 1***. A list of abbreviations is listed in ***Supp. Table 1***.

**Table 1.**
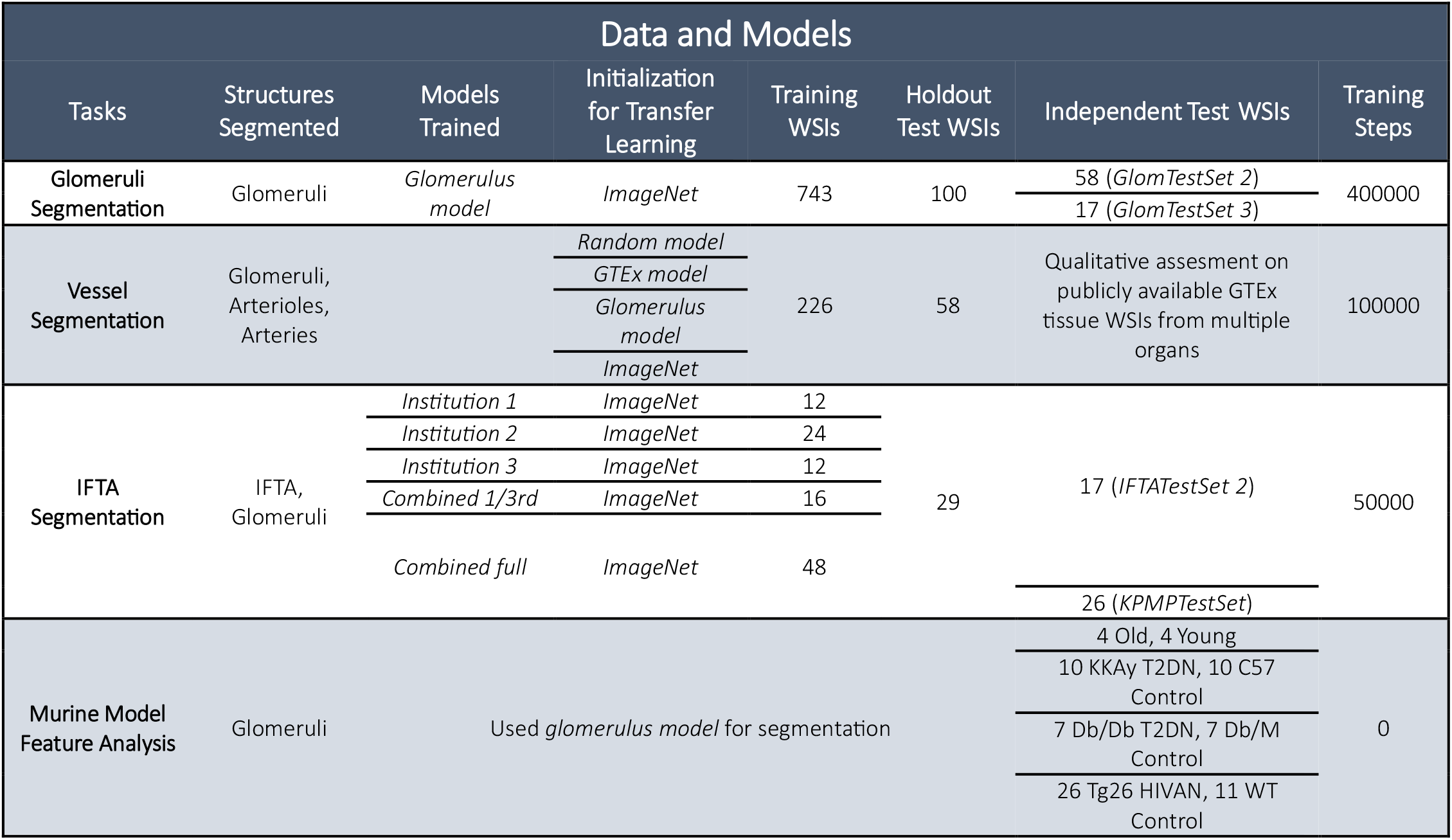
Data used and models trained. Different segmentation tasks, corresponding trained models, segmented structures, initial model used for transfer learning, WSIs used for training, hold-out testing, and independent testing, and training steps. We note that *GlomTestSet 2* is the same as the Vessel Segmentation holdout dataset (58 WSIs). *GlomTestSet 3* is also the same as *IFTATestSet 2* (17 WSIs).

### HISTO-CLOUD

Using the simple cloud-based interface, users can upload WSIs and train a segmentation network using their own annotations (see ***Fig. 1b)***. Users can iteratively apply *Histo-Cloud*’s training and prediction plugins in an active learning framework, to build up powerful segmentation models with reduced effort^7^. The segmentations produced by *Histo-Cloud* are converted to contours or heatmaps for direct display on the WSIs. When developing new segmentation models, the slide-viewing environment of this tool, enables rapid qualitative evaluation of algorithm progress by displaying the network predictions (***Fig. 1a***).

**Fig. 1.**
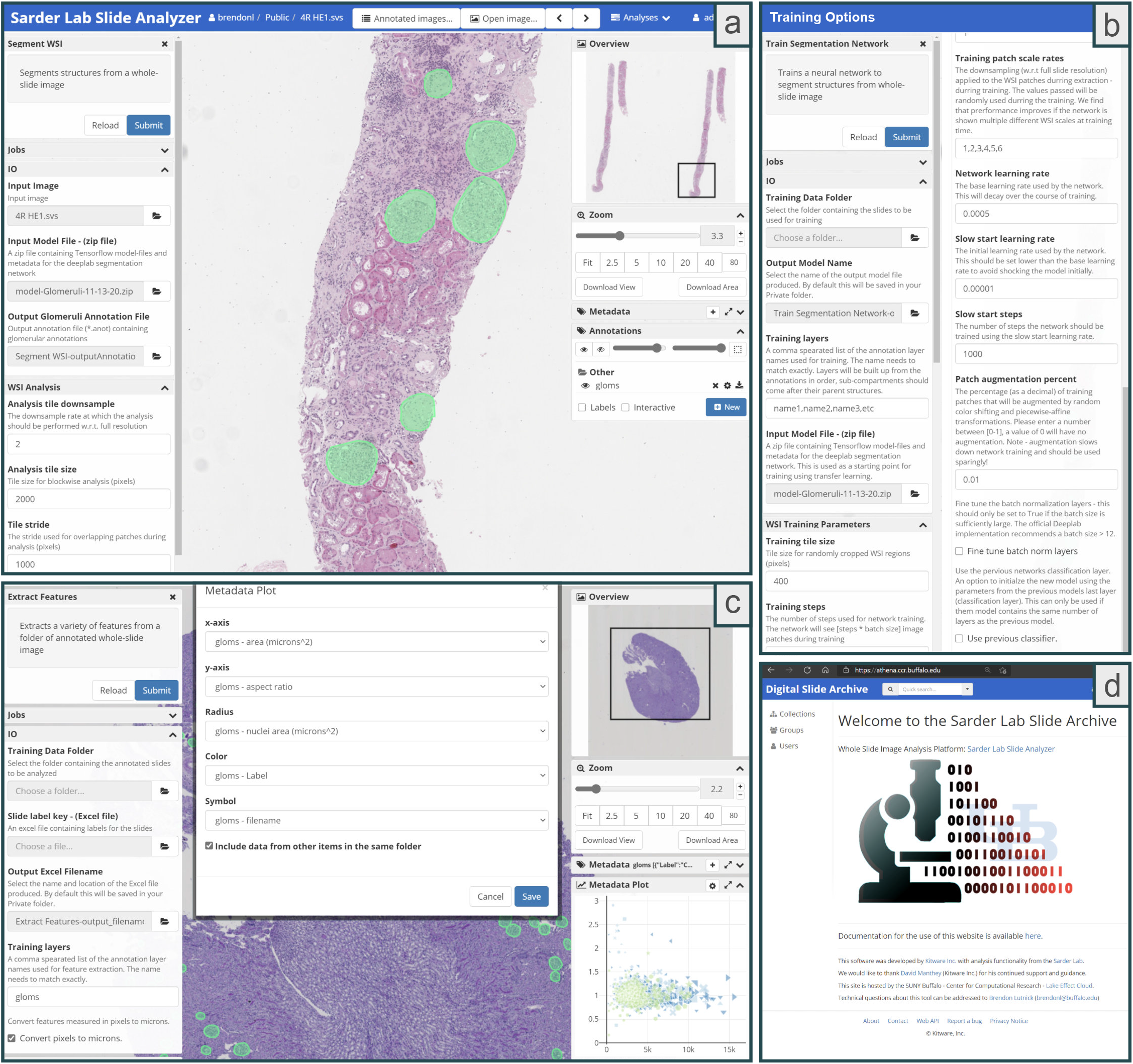
The user interface of the segmentation tool (available via the web). **[a]** The left *<Segment WSI>* column shows the controls for the segmentation plugin: *<IO>* is required arguments and *<WSI Analysis>* contains optional parameters. The right column shows the WSI viewer controls and annotations created by the plugin. The green annotations on the holdout slide are predicted by the plugin and are easily editable by the user. Slides are analyzed by clicking the *<Submit>* button in the top left corner, which submits a segmentation job, running the DeepLab network on the remote server (where HistomicsUI is installed). **[b]** The options from the *<Train Segmentation Network>* plugin. Under the *<IO>* section, a user can specify a directory full of annotated WSIs to use for network training with the *<Training Data Folder>* option, and where to save the trained model with the *<Output Model Name>* option. The *<Training layers>* option gives users the ability to choose which of the annotation layers should be used for training, and subsequently single or multi-class segmentation models can be trained. To speed up the training process, a previously trained segmentation model can be used for transfer learning by specifying the *<Input Model File>*. Hyper-parameters for training the network can be specified using the options in the *<WSI Training Parameters>* section – the values herein set to defaults which we have found work well. **[c]** shows the *<Extract Features>* plugin which can be used to extract image and morphology features from annotated objects. These features are written to the slide metadata and can be plotted from within the online interface via the *<Metadata Plot>* tab (on the right). **[d]** shows the welcome screen of the online interface athena.ccr.buffalo.edu.

Going beyond segmentation, an included modular plugin extracts features from segmented WSI tissue regions. These features are written into the metadata of uploaded slides and can be exported in spreadsheet form for further analysis. We have included a plotting tool in the user interface of the online slide viewer for quick exploration of these extracted features, ***Fig. 1c***.

The source code can be run traditionally via the command line, but we expect the majority of users will utilize the intuitive HistomicsUI based cloud interface. The source code is available on GitHub at https://github.com/SarderLab/Histo-cloud, and packaged as a pre-built Docker image^14^ https://hub.docker.com/r/sarderlab/histo-cloud. This data sharing allows for easy deployment on a remote server for use as well as further development by the community over the web. Additionally, a publicly available instance of *Histo-Cloud* is available for the community at: athena.ccr.buffalo.edu. All the models described are available in the *<Collections>* section in the *<Segmentation models>* folder on athena.ccr.buffalo.edu or at https://bit.ly/3ejZhab. Documentation for using this tool is available at https://bit.ly/3nNMpfH.A video overview of *Histo-Cloud* is available at https://bit.ly/3r5GrZr.

### GLOMERULAR SEGMENTATION – scalability

To assess the computational *scalability* of *Histo-Cloud*, a network model for glomeruli segmentation was trained using renal tissue WSIs (*glomerulus model*). This network was trained using a very large dataset containing 743 WSIs (*GlomTrainSet*). Network performance was evaluated on a holdout set of 100 additional human renal tissue WSIs (*GlomTestSet 1*). The computationally generated segmentation was robust when compared with manual annotations for glomeruli and generated the following statistics: *F-score=0.97, Matthews correlation coefficient* (*MCC*)*=0.97, Cohen’s kappa=0.97, intersection over union* (*IoU*)*=0.94, sensitivity=0.95, specificity=1*.*0, precision=0.99, and accuracy=1*.*0*. This model also performed robustly on two independent test WSI datasets (*GlomTestSet 2 & 3*) originating from an institution not included in the training dataset with ground-truth established by a separate annotator (*MCC=0.83* and *0.90* on *GlomTestSets 2* and *3*, respectively) (***Fig. 2a***). ***Fig. 2c*** shows examples of glomerulus segmentation performance for a diverse set of glomerular pathologic changes and histochemical stains.

**Fig. 2.**
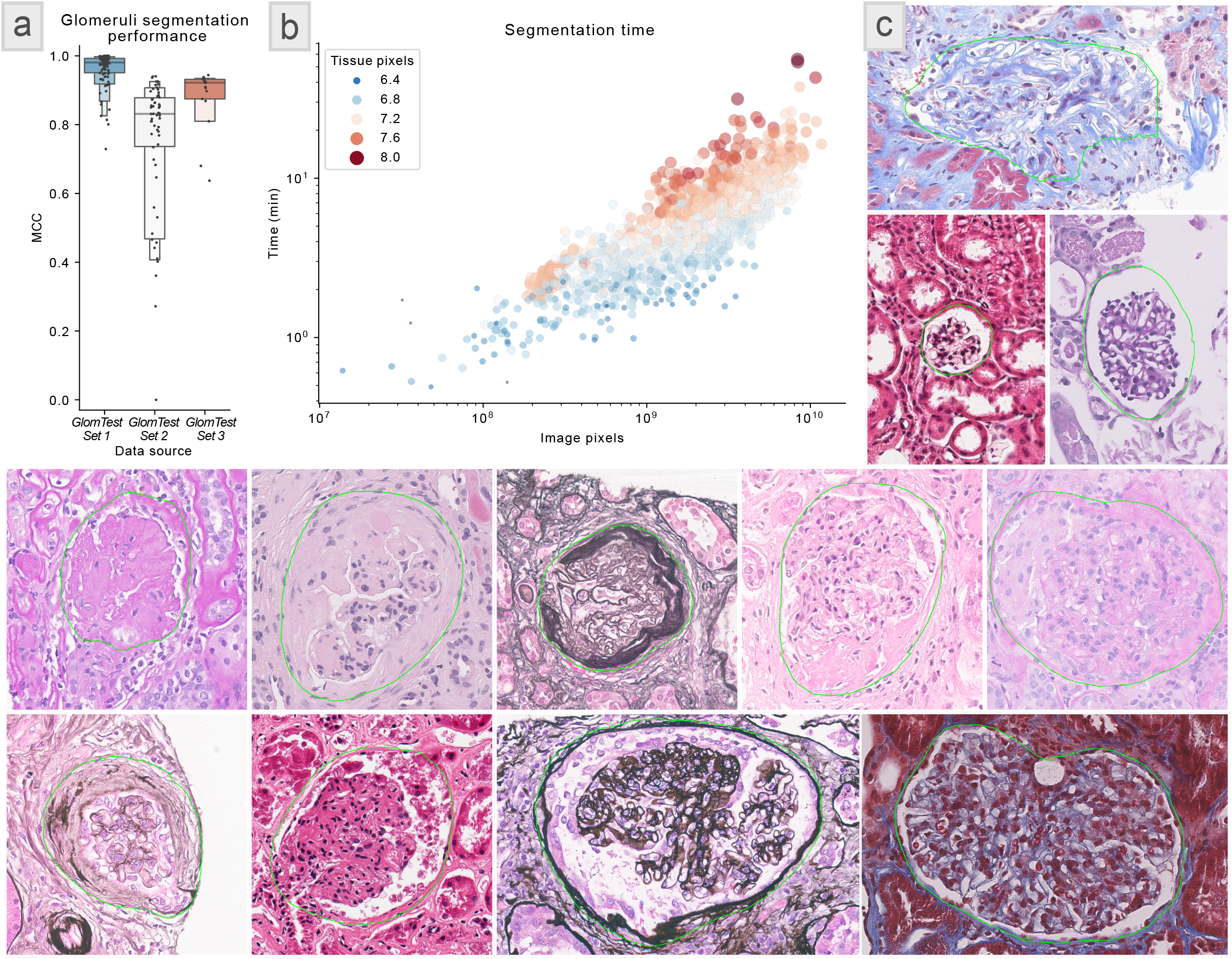
Glomeruli segmentation results – scalability study. **[a]** The segmentation performance of *glomerulus model* for glomeruli detection. Mathews correlation coefficients were calculated for three renal tissue WSI datasets, as specified in subsection *GLOMERULI SEGMENTATION – scalability* under the section Results: *GlomTestSet 1* contained 100 WSIs holdout from the training set *GlomTrainSet, GlomTestSet 2* had 58 WSIs, and *GlomTestSet 3* had 17 WSIs. Both *GlomTestSet 2* and *GlomTestSet 3* were from an institution independent of the institutions from where the training dataset *GlomTrainSet* was formed for training the *glomerulus model*. Further, glomerular boundaries in *GlomTestSet 2* and *GlomTestSet 3* were annotated by an independent annotator who was not involved in annotating glomeruli in *GlomTrainSet*. **[b]** shows the prediction time in minutes as a function of the WSI size in pixels for glomeruli predictions on 1528 WSIs in *GlomTestSet 4*. The color and size of the points represent the size of the automatically extracted tissue region of the slide (the analyzed region) in pixels. The proposed glomerular segmentation model scales roughly linearly in time for increasing WSI size. **[c]** A batch of randomly selected glomeruli with the computationally segmented boundaries from the 100 holdout WSIs in *GlomTestSet 1*. This selection is intended to highlight the diversity of pathology and staining of the holdout dataset.

We have found the performance of *Histo-Cloud* continually improves while achieving high specificity when deployed in a human in the loop setting. This process allows experts to iteratively correct the network predictions on holdout WSIs before incorporating them into the training-set, and the subsequent training reduces future annotation burden^7^. This process is facilitated due to the ability of our system to view predictions interactively on the WSIs via the web interface, which is helpful to determine WSIs where the trained model struggles. We used this strategy to train the *glomerulus model* iteratively and obtained a decreasing number of incorrect segmentations with increasing iterations.

As part of the scalability study the segmentation speed was assessed. Prediction time as a function of WSI size was tracked on a set of 1528 WSIs (median time=4.7 min, median size=1.9 Gigapixels) from a set that have similar diversity as in *GlomTrainSet*, we refer to this set as *GlomTestSet 4. Histo-Cloud* uses hardware acceleration on the host server to speed processing and is capable of segmenting a large histology section in as little as 1 min. The segmentation time depends (approximately linearly) on the size of the tissue section; ***Fig. 2b*** quantifies segmentation speed as a function of image pixels on WSIs from *GlomTestSet 4*. The algorithm performs a fast thresholding of the tissue region within the slide to reduce the computational burden for slides with large non-tissue areas. There is a slight programmatical overhead when opening and caching larger slides, this appears as a gentle upslope of points of the same color in ***Fig 2b***.

### VESSEL SEGMENTATION – adaptability

To evaluate the *adaptability* of *Histo-Cloud* for segmenting multiple structures from WSIs, we retrained the *glomerulus model* to segment glomeruli, arteries, and arterioles. The training set is referred to as *VessTrainSet*, and the test set is *VessTestSet*.

Transfer learning is a machine learning technique where a model developed for one purpose is retrained for another purpose^15^. Using the *glomerulus model* as the starting point for transfer learning, *Cohen’s kappa* of 0.86 was obtained for segmenting arteries, arterioles, and glomeruli. The *kappa* metric was computed based on pixelwise agreement between computational segmentation and manual ground-truth. To study the effect of transfer learning on segmentation performance, we trained another model by randomly initializing the network parameters (*random model*); performance decreased to *Cohen’s kappa* of 0.51.

We further explored the possibility of improving the computational performance without access to a model trained from a large segmented dataset. Toward this goal, we used the Genotype-Tissue Expression dataset (GTEx)^16^, which contains 15989 H&E stained WSIs from 40 different tissue types, to pre-train a segmentation model to detect the tissue type. This was accomplished without any human annotation, by thresholding the tissue region of each slide and training a model to classify the tissue type of each slide. The goal was to create a model for transfer learning which had been exposed to diverse tissue morphologies, and therefore had learned filters useful for more fine-grained segmentation tasks. While transfer learning using the resulting model (*GTEx model*) did improve the segmentation performance of glomeruli, arteries, and arterioles (*Cohen’s kappa=*0.68) over random initialization, performance was below that achieved using the *glomerulus model*.

Finally, we trained a fourth model, transfer learning with a model pretrained on the *ImageNet*^17^ dataset, this same model was originally used to train the *glomerulus model*. Surprisingly, this model (*ImageNet model*) achieved the best segmentation performance (*Cohen’s kappa=0.87* with *MCC* of *0.86,0.66*, and *0.91* for segmenting arteries, arterioles, and glomeruli, respectively). This performance improvement is further explored in the *DISCUSSION* section. A more detailed comparison of these results is shown in ***Fig. 3a***, with randomly selected holdout predictions from *VessTestSet* in ***Fig. 3b***. To explore the performance of the *ImageNet model* on an independent test set, we segmented GTEx WSIs from different organs, examples are shown in ***Fig. 3c***.

**Fig. 3.**
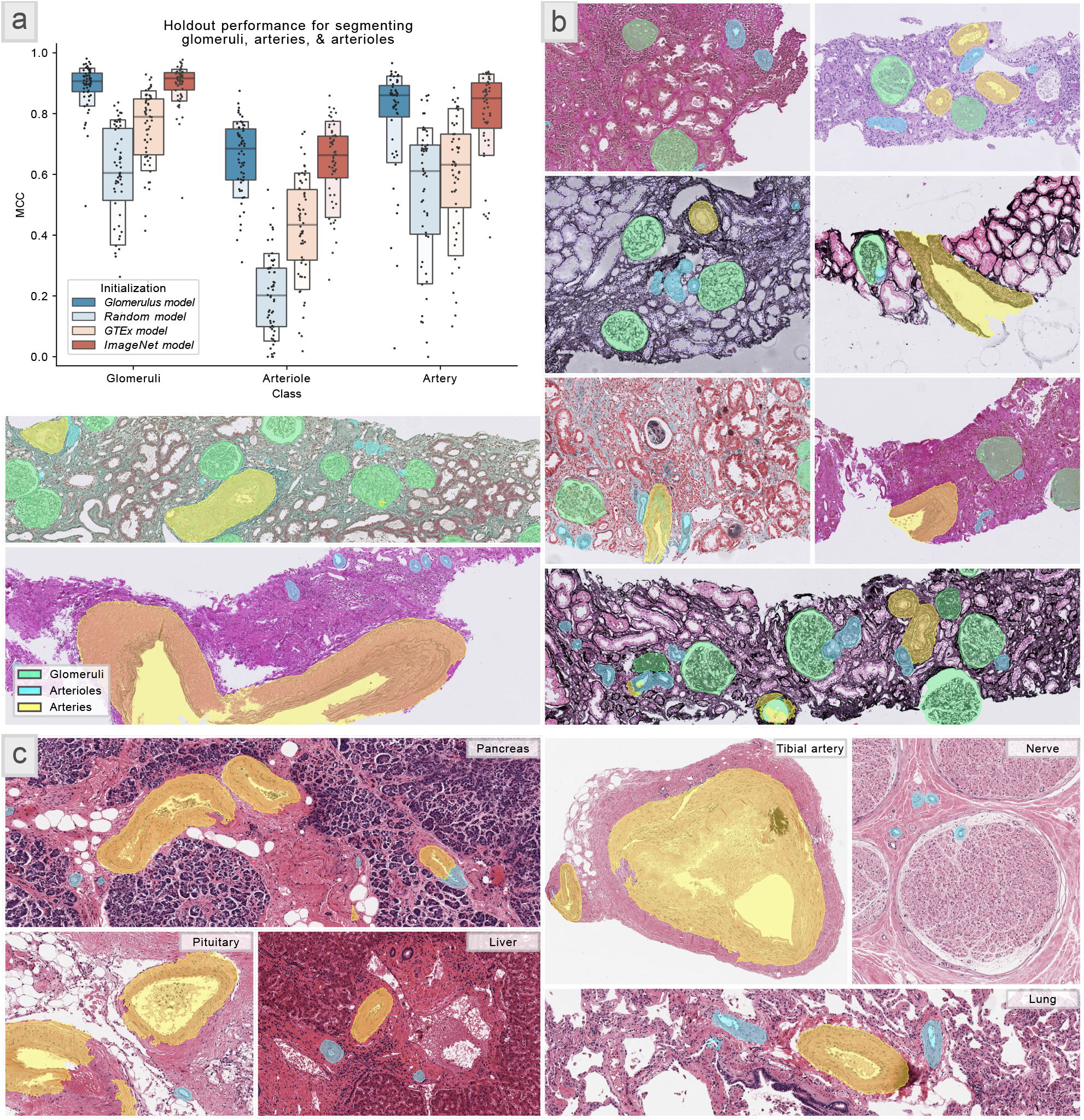
Vessel segmentation results – transfer learning study. **[a]** The segmentation performance as a function of network initialization (measured as Matthews Correlation Coefficient) for the *VessTestSet* with 58 holdout WSIs from the WSIs used for vessel segmentation. Note that glomeruli were also segmented here. The ground-truth annotations of structures were generated for three classes: glomeruli, arterioles, and arteries. The colors here represent different transfer learning sources for parameter initialization. Namely, ***glomerulus model*** is the model used for glomerular segmentation results in ***Fig. 2***, and results in a *Cohen’s kappa=0.86* here when this model was used for initializing the training. ***Random model*** does not use transfer learning for parameter initialization, and results in a *Cohen’s kappa=0.51* when used as the initialization strategy for training. ***GTEx model*** is a model trained to identify the diverse tissue types from the publicly available GTEx tissue WSI dataset (15989 WSIs with 40 different tissue types), and results in a *Cohen’s kappa=0.68*, when used for training initialization. ***ImageNet model*** uses a model pre-trained on the *ImageNet* dataset, and produces a *Cohen’s kappa=0.87*, when used for initializing the training. Performance for computationally segmenting three classes, namely, glomeruli, arteries, and arterioles, with respect to the ground-truth was measured using Cohen’s kappa metric based on pixelwise matching between computational segmentations and ground-truth. **[b]** shows randomly selected crops of WSIs from the holdout set (*VessTestSet*) with computational segmentations by the model trained based on *ImageNet* model as the starting point. **[c]** shows randomly selected crops of various types of tissues from GTEx WSIs, computationally segmented using the model trained based on *ImageNet* model as the starting point. Despite being trained only on kidney tissues, the trained model is able to segment arteries and arterioles in diverse tissue types. We also note that the GTEx slides are autopsy tissues scanned at 20X, and the training set for this study *VessTrainSet* was scanned at 40X, and did not contain autopsy tissue WSIs.

### INTERSTITIAL FIBROSIS & TUBULAR ATROPHY (IFTA) SEGMENTATION – adaptability

To further evaluate the *adaptability* of *Histo-Cloud*, the effect of dataset variability on the segmentation of IFTA was studied in a distributed setting; namely, our web-based setup (in cloud). IFTA is morphological changes in the renal cortex reflecting “chronic” injury with resultant scar formation and is an important indicator to predict renal disease prognosis^9^.

To generate a ground truth, three pathologists provided WSIs from their institutions and manually annotated IFTA. Past studies have shown significant disagreement among pathologists in manually annotating IFTA^9^. To minimize such disagreement, the pathologists used the definition of IFTA based on Banff 2018 criteria^18^, and also collaborated via our web-based tool in a distributed setup for IFTA annotation. Further the inclusion criteria of cases (discussed in the *Methods - WSIs from IFTASet 1, IFTASet 2, IFTASet 3, IFTATestSet 2, and GlomTestSet3* section) minimized variability of the annotation process.

A holdout dataset was randomly selected by pooling one-third of the slides from each institution *(n=29)*. We refer to this set as *IFTATestSet 1*. Another dataset from a fourth institution (*IFTATestSet 2, n=17*) was used for independent testing. A pathologist from this fourth institution manually annotated IFTA in *IFTATestSet 2* to generate the ground-truth.

We trained five models for IFTA segmentation using the pathologist provided ground-truth: the first three models were trained using slides from a single institution - *IFTASet 1* (12 slides), *IFTASet 2* (24 slides), and *IFTASet 3* (12 slides). We refer these as *Institution 1, 2, & 3* models respectively. The fourth model used the combined training data from all the three sets (48 slides), referred to as *Combined full*. A final model used 1/3^rd^ of this combined set (16 WSIs), ensuring the amount of training data was comparable to the first three models. This model is referred as *Combined 1/3*^*rd*^.

To better assess the performance of the trained models, we output the network logits (predictions prior to using the argmax function) which were used to construct ROC plots for each model. This process allowed us to display IFTA predictions as heatmaps in HistomicsUI (***Fig. 4d***). Interestingly on *IFTATestSet 1* training with 1/3^rd^ of the combined dataset (*Combined 1/3*^*rd*^ model) yielded better IFTA segmentation (*AUC=0.93*) than training with a single institution dataset alone (***Fig. 4a***) (*AUC=0.78,0.76, and0.91* for models *Institutions 1, 2*, & *3*, respectively). When we tested the *Combined full* model, the performance improved to *AUC=0.95*. The same trend was observed when segmenting IFTA in the independent test set *IFTATestSet 2* (***Fig. 4b***), with *AUC=0.68,0.75, and0.83* for models *Institution 1, 2, & 3*, respectively, *AUC=0.86* for *Combined 1/3*^*rd*^ model, and *AUC=0.88* for *Combined full* model.

**Fig. 4.**
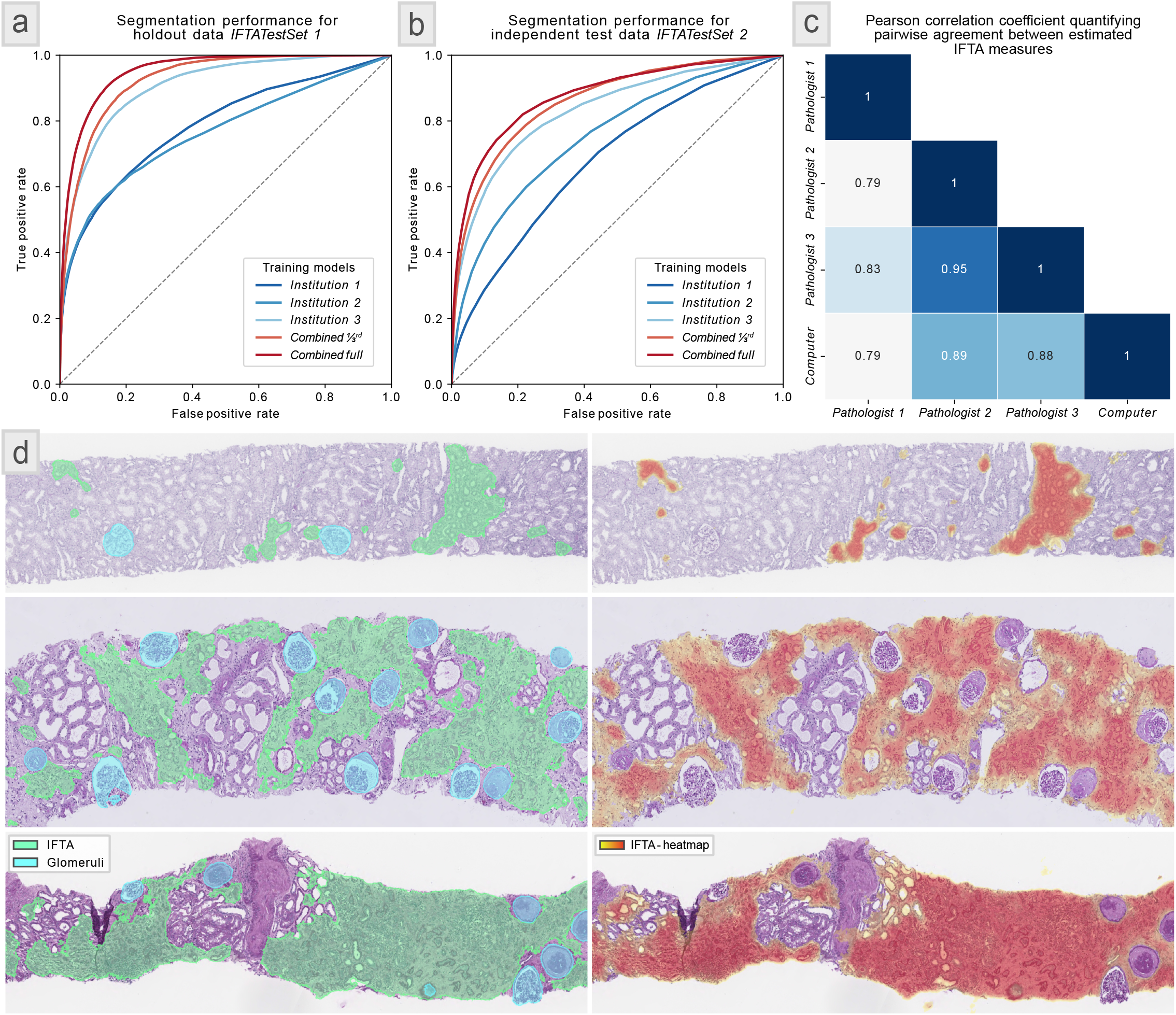
IFTA segmentation results – multi-institute study. **[a]** ROC plots showing the segmentation performance of the five trained IFTA models on the 29 holdout WSIs, *IFTATestSet 1*. The models (*Institution 1, Institution 2*, and *Institution 3*) were trained using training datasets from three different institutions, namely, *IFTASet 1* with 12 WSIs, *IFTASet 2* with 24 WSIs, and *IFTASet 3* with 12 WSIs. These datasets were annotated by three respective pathologist annotators, marking the ground-truth IFTA boundaries. The *Combined full* model was trained using all the 48 WSIs from all the three datasets. The *Combined 1/3*^*rd*^ model used 1/3^rd^ of the pooled training set, randomly selected from the three datasets (16 WSIs), to have comparable number of training WSIs as used by the first three training datasets. This last model yielded a better performance in segmenting IFTA than the models, *Institution 1, Institution 2*, and *Institution 3*, suggesting that a more diverse dataset improves segmentation performance. *Combined full* model offered slightly better performance than *Combined 1/3*^*rd*^ model. **[b]** shows the performance of the five models using ROC measures on the independent test dataset *IFTATestSet 2* with 17 WSIs. This dataset was originated from an institution independent than those sourced for results shown in **[a]**, and was annotated by an independent annotator than those employed for **[a]**. Using *IFTATestSet 2*, we observed the same performance trend as in **[a]. [c]** shows the pairwise Pearson correlation coefficients (*p*-value < 0.05) for percent IFTA scored visually manually by three additional annotators and the percent IFTA estimated based on computational segmentation using the *Combined full* model (computer) for the 26 CKD renal tissue biopsy WSIs in *KPMPTestSet*. The KPMP cohort acted as another independent test set which was never seen by our trained model. **[d]** shows computational IFTA predictions using the *Combined full* model on the holdout WSIs *IFTATestSet 1*. The left shows the traditional contour predictions, the right shows the corresponding heatmap predictions developed specifically for structures with poorly defined boundaries.

The IFTA segmentation models were trained to simultaneously segment IFTA and glomeruli. We observed the same performance trend for glomerulus segmentation via the IFTA models in both *IFTATestSet 1* and *2;* these results are available in ***Supp. Fig. 2***. The ROC plots (generated by thresholding the network logits) for all the glomeruli, artery, and arteriole segmentations conducted in this work are shown in ***Supp. Fig. 3***.

To demonstrate the robustness in another independent cohort and compare the trained model to visual manual estimation of IFTA done in the clinical setting, we used an additional 26 PAS-stained chronic kidney disease renal biopsy cases from the Kidney Precision Medicine Project (KPMP)^19^ consortium. We refer this set as *KPMPTestSet*. Three KPMP pathologists, provided a percent IFTA score to the nearest 10 percent for each slide following Banff 2018 definitions^18^. This scoring was done via visual estimation, without any annotation on the slides. The five IFTA segmentation models discussed above were used to segment IFTA boundaries in the *KPMPTestSet*, percent IFTA was estimated as segmented IFTA area over total renal cortex area, and the resulting computationally estimated scores were correlated with the manual visual estimation. ***Fig. 4c*** shows a confusion matrix describing correlations (*p*-value < 0.05) between pathologists and the computer for the *Combined full* model. We found that the correlation measures among pathologists and the computer models were comparable. ***Supp. Fig. 4*** shows a full comparison of the five IFTA segmentation models and each KPMP pathologist. ***Fig. 4d*** depicts examples of qualitative IFTA segmentation performance.

### MURINE MODEL ANALYSIS – utility

Finally, we show the *utility* of *Histo-Cloud* in a basic research application, analyzing digital image features extracted from computationally segmented glomeruli (via the G*lomerulus model*) from four murine models. A description of the models used is available in *Methods* - *WSI from murine kidney tissue*.

WSI from each model contained multiple sections obtained from one murine, with an average of 90-200 glomeruli per section. For the current analysis, we extracted 315 engineered image features from each segmented glomerulus. Feature definitions and quantification methods are discussed in our prior work^5^, a description is also available in ***Supp. Table 2***. The features were selected to reflect active, present, and physical manifestations of kidney pathophysiology. We used an unsupervised Uniform Manifold Approximation and Projection (UMAP)^20^ to learn a two-dimensional manifold in the feature space (performing dimensionality reduction). Each glomerulus was plotted (with label) in this space to visualize the separability between classes (control vs disease) in each murine model (***Fig. 5***). To quantify this separability, we trained a *K*-nearest neighbor (KNN)^21^ classifier using the UMAP features with five-fold cross validation, and computed the optimal *Cohen’s kappa* achieved over multiple *K* for each murine model (***Supp. Fig. 5***). Overall, we found the aging, KKAy and Db/Db diabetes models to have good unsupervised class separability (***Figs. 5a, b, c***). We also applied Seurat^22^ software to analyze the image feature data and to characterize differential feature abundance. The distribution of the top feature separating control from disease, and the most representative glomeruli image patches depicting differences from these two classes are shown in ***Fig. 5***.

**Fig. 5.**
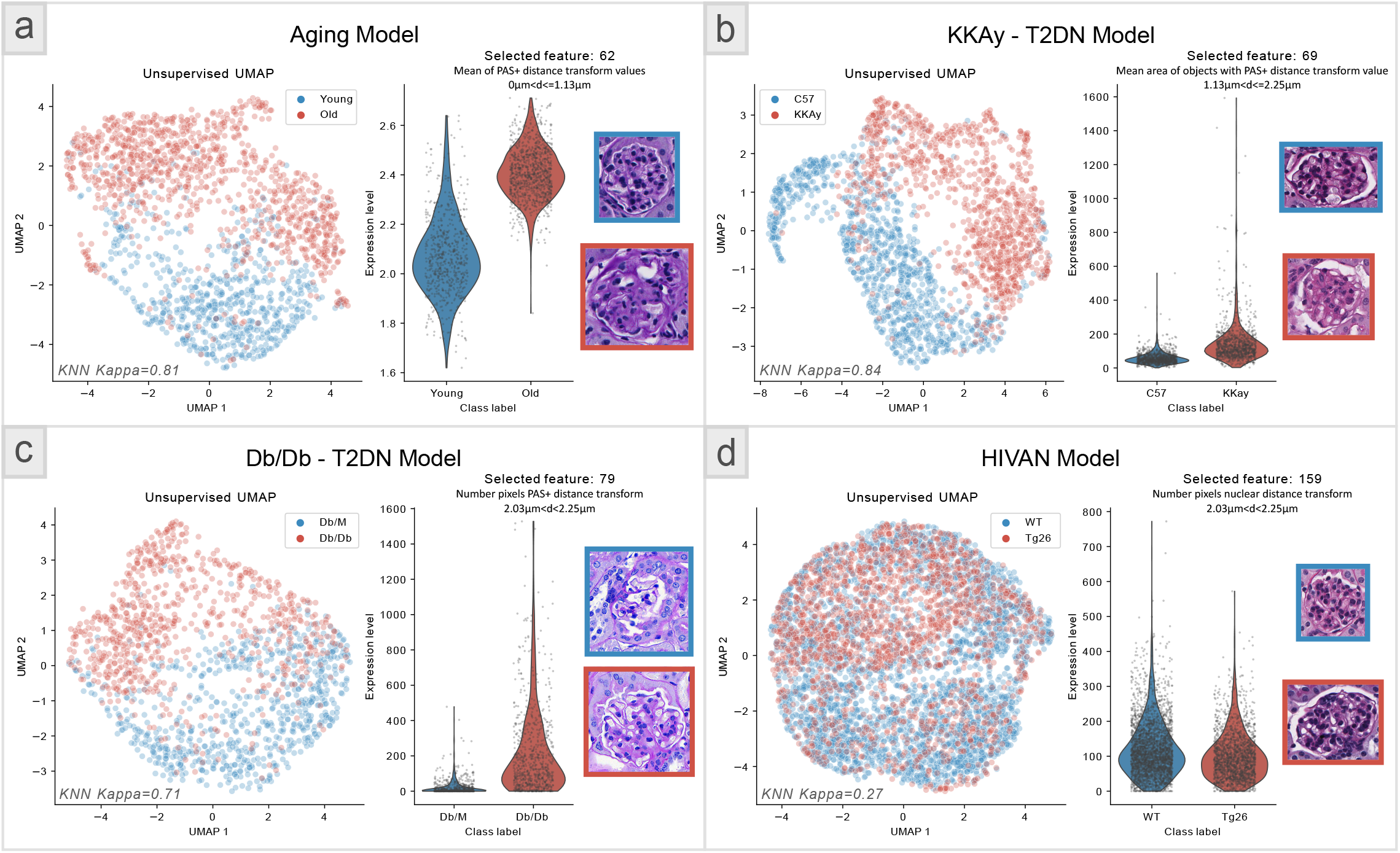
Murine model glomerulus feature analysis – utility study. Feature analysis from glomeruli segmented from renal tissue WSIs from four murine models: **[a]** is an aging model, **[b & c]** are two type 2 DN models (KKAy and Db/Db), and **[d]** is an HIVAN model. In each panel the left plot shows an unsupervised UMAP representations of 315 engineered image features extracted from the murine glomeruli, where the glomeruli were segmented using the *glomerulus model*. Here each dot is a glomerulus and the red and blue colors differentiate the disease from control. Definitions and quantification strategy of the 315 engineered image features are available in our prior work^5^. Separation between the disease and control classes are quantified using *KNN kappa* metric. The right plot shows the highest differentially expressed feature as predicted using the Seurat software^22^. The representative glomeruli from each murine class depicting this differentially expressed feature, and the feature value, are shown on the right for each murine model. Definitions of the 315 features are provided in ***Supp. Table 2***. This study suggests that the seamless segmentation of glomeruli from large WSIs using our tool facilitates conducting deep glomerular feature analysis to study novel murine models.

## DISCUSSION

In this work, we contribute three elements to the digital pathology community to advance tissue analysis: an online tool, the source code, and trained segmentation networks. We believe that easy-to-use AI tools and collaborative development of powerful models will benefit the digital pathology research community.

This work was motivated by our previously developed Human-AI-Loop (H-AI-L)^7^ which allows for iterative annotation of WSIs significantly reducing the annotation burden. As most work in computational pathology, H-AI-L has found limited utilization by the pathology research community due to complexities of installation. To address this limitation, we implemented *Histo-Cloud* as an online tool which does not require the installation of any software on the user’s local computer. All processing occurs on the remote server, which hosts the web client. Like the original H-AI-L work, we use the DeepLab segmentation network^23^ for processing image patches, but *Histo-Cloud* uses on-the-fly processing of WSI patches to increase the tool’s performance and scalability. Data permissions (set via the digital slide archive - DSA^13^) can be adjusted to keep uploaded data secure.

Annotation done interactively on the WSI fits easily into pathologist workflow, and the cloud-based nature of *Histo-Cloud* abstracts any computational overhead away from the end-user. Annotation can be done on any internet connected device without any software installation. lf the user prefers to annotate locally, we have added options to ingest and export annotations in an Extensible Markup Language (*XML)*^24^ format readable by the commonly used WSI viewer Aperio ImageScope^25^. The authors note two complimentary works: HistomicsML^26^ and Quick Annotator^27^, both use superpixels^28^ and active learning^29^ to speed the annotation process. HistomicsML also uses HistomicsUI for deployment, and Quick Annotator is run locally in the QuPath slide viewer^30^. A future extension of our tool will combine edge detection and snapping^31^ to speed the initial segmentation by human annotators.

Conducting the transfer learning study using the GTEx tissue histology WSIs (***Fig. 3a***) (15989 WSIs containing 2.6 trillion total image pixels, 4.7 TB of data) and training the *glomerulus model* for glomeruli segmentation (***Fig. 2a***) (743 WSIs, 1.8 trillion pixels, 276 GB) were stress tests for scalability. Setting *Histo-Cloud*’s accessibility benefits aside, the study of glomeruli segmentation (***Fig. 2***) not only uses the largest most-diverse cohort of WSIs, but also reports the best performance in the literature for glomerular segmentation. In our previous work on H-AI-L^7^ we trained Deeplab-v2^32^ using a dataset of 13 PAS and hematoxylin and eosin (H&E) stained murine WSIs containing 913 glomeruli, and achieved an *F-score=0.92. Kannan et al*.^33^ used Inception-V3^34^ for the sliding window classification of glomeruli with a set of 885 patches from 275 trichrome stained biopsies, and reported *MCC=0.63. Bueno et al*.^35^ trained U-net^6^ with 47 PAS stained WSIs, and reported *Accuracy=0.98. Gadermayr et al*.^*36*^ used 24 PAS-stained murine WSIs to train U-net^6^, reporting *Precision=0.97* and *Sensitivity=0.86*.

*Jayapandian el al*.^*37*^ present the most comprehensive results on glomeruli segmentation, training U-net^6^ on a dataset containing 1196 glomeruli from 459 human WSIs stained with H&E, PAS, Silver and Trichrome, reporting *F-Score*=0.94. However their analysis is limited to glomeruli with minimal change disease^38^. In contrast our training dataset *(GlomTrainSet*) contained 743 WSIs from both human and mice, stained with diverse histological stains, with a 61734 total glomeruli, from diverse disease pathologies beyond minimal glomerular changes. The holdout dataset *GlomTestSet 1* contained similar diversity (***Fig. 2c***). Our trained model also performed well on independent test datasets *GlomTestSet 2 & 3* (***Figs. 2a***). Predictably, performance on the independent test datasets *GlomTestSet 2 & 3* was lower than the holdout dataset, albeit visual assessment of the segmentations appeared consistent with expert opinion. The modularity *of Histo-Cloud* will allow others to adapt the trained model to include more structurally abnormal glomeruli.

When testing the effectiveness of transfer learning, we found that adapting the *ImageNetmodel* for segmenting glomeruli, arteries, and arterioles using the *VesselTrainSet*, performed better than using the *glomerulus model* as the starting point (albeit marginally). The *ImageNet model*was trained on thousands of natural image classes, and is widely used in computer vision literature^17^. It is surprising that despite having seen renal tissue the *glomerulus model* reduced feature generalizability. This result suggests that it is a better practice to start network training using the *ImageNet* parameters (now the default for training *Histo-Cloud* models in the cloud). Encouragingly, when applying the developed vessel segmentation model to different tissue types from the publicly available GTEx tissue WSIs^16^, the segmentation of arteries and arterioles was found to be consistent with expert opinion (***Fig. 3c***).

Perhaps the most interesting aspect of a cloud-based segmentation tool is the ease of crowd sourcing annotation and developing collaborative models across centers or institutions^39^. As discussed above and also known that manual annotation of IFTA boundaries by multiple pathologists suffer from high degree of disagreement^9^. In contrast, *Histo-Cloud’s* web-based system allowed the annotators to view each other’s annotations in annotating *IFTASet 1, 2*, & *3*, and *IFTATestSet 2* for the multi-institute IFTA study (see *IFTA SEGMENTATION – adaptability* under *RESULTS*). We further note that visualizing IFTA prediction confidence using heatmaps was more reflective of the underlying biology than using contours, confirmed by subject matter experts via visual assessment. Namely, a heatmap depicts a probability which is more informative than contours which display binary predictions. Examples of IFTA segmentations on the holdout data *IFTATestSet 2* as both contours and heatmaps are shown in ***Fig. 4d***. The functionality to output segmented regions as heatmaps is available using the segmentation plugin.

The IFTA segmentation study further highlights the importance of training set diversity. Training using data from more institutions improved segmentation performance, even when less WSIs from each institution were used. Namely the performance of *Combined 1/3*^*rd*^ *model* in comparison to *Institution 1, Institution 2*, and *Institution 3* models (see *IFTA SEGMENTATION – adaptability* under *RESULTS*). This and the results described in the previous paragraph suggest a cloud-based environment is ideal for the development of models for histology segmentation, avoiding bias and allowing easy interaction between annotators for generating ground-truth by centralizing data from multiple institutions. Users can choose to pool their data or simply utilize models trained by others to aid in annotation or for transfer learning.

Finally, the murine model analysis case study suggests that our tool will enable basic science laboratories working on murine experiments to study differential abundant image features in various disease models as well as in treatment groups. In summary, the analytic approaches described here will enable researchers who lack software engineering skills to analyze the histopathology from murine models or human tissue, using an intuitive online cloud-based framework.

## METHODS

### WSIs for GlomTrainSet, GlomTestSet 1, and GlomTestSet 4

These datasets were used for the segmentation of glomeruli. This dataset consists of both human and murine renal tissue WSIs from various institutes as well as publicly available repositories, using diverse stains and different scanners. The institutions included the University of California at Davis (UC Davis), Johns Hopkins University (JHU), Kidney Translational Research Center (KTRC) at Washington University School of Medicine at St. Louis (WUSTL), Seoul National University Hospital Human Biobank (SNUHHB), Vanderbilt University Medical Center (VUMC), University at Buffalo (UB), University Hospital Cologne (UHC), and the publicly available Genotype-Tissue Expression (GTEx) portal, a repository that hosts human autopsy WSIs.

The *GlomTrainSet* consisted of 743 WSIs, 428 from human and 315 from murine tissues, containing a total of 61734 manually verified glomerular annotations. *GlomTestSet 1* consisted of 100 holdout slides from the same data sources as *GlomTrainSet*. This included 3816 glomeruli, 37.8 GB of compressed image data, and a combined total of more than 0.24 trillion image pixels. *GlomTestSet 4* contained an additional 1528 WSIs from the same sources that were used to study the scalability and prediction time of the method.

The human renal tissues manifest disease pathology spanning various stages of diabetic nephropathy; various classes of lupus nephritis; renal transplant protocol biopsies, including time-zero, protocol, and indication biopsy cases; human autopsy renal tissues publicly available via GTEx with diversity in age, sex, and race; and renal biopsies with pathologies that include membranous nephropathy, thrombotic microangiopathy, pauci-immune glomerulonephritis, focal segmental glomerulosclerosis (FSGS), mesangiopathic glomerulonephritis, arteriolosclerosis, hypertension, IgA nephropathy, chronic tubulointerstitial nephritis, acute tubular necrosis, Fabry disease, amyloid nephropathy, membranoproliferative glomerulonephritis, light chain cast nephropathy, minimal change disease, post-infectious glomerulonephritis, idiopathic nodular glomerulosclerosis, and anti-glomerular basement membrane disease. The human data was collected in accordance with protocols approved by Institutional Review Board at the UC Davis, JHU, KTRC, WUSTL, SNUHHB, VUMC, and UB. The SNUHHB data was shared under IRB number H-1812-159-998.

Murine renal tissues included in *GlomTrainSet* and *GlomTestSet 1* came from three different models. For the first model wild-type FVB/N mice were subjected to a combination of four interventions that induce a post-adaptive form of FSGS. The interventional process includes 0.9% saline drinking water, angiotensin II infused via osmotic pump, uni-nephrectomy, and deoxycorticosterone delivered by implantation of a subcutaneous pellet, summarized as the SAND model^40,41^. The second model was a streptozotocin (STZ) diabetes murine model that manifests nephropathy; a detailed description of this model is discussed in our prior work^42^. The third model was a nephrin knockdown (nephrin KD) murine model, was implemented using a published protocol^43^, and shows mesangial hypercellularity and sclerosis, glomerular basement membrane thickening, and podocyte loss.

The tissues were sectioned at 2-5 µm thickness for staining and imaging. The data consist of tissues stained with diverse histological stains, including hematoxylin & eosin (H&E), periodic-acid Schiff (PAS) with hematoxylin (PAS-H) counterstain, Silver, Trichrome, Verhoeff’s Van Gieson, Jones, and Congo red. The slides were scanned using different brightfield microscopy WSI scanners, including Aperio VERSA digital whole slide scanner (Leica Biosystems, Buffalo Grove, IL), Nanozoomer (Hamamatsu, Shizuoka, Japan), and MoticEasyScan Pro (Motic, San Antonio, TX), at 40X resolution. Pixel resolution of the images used was 0.13 µm to 0.25 µm.

### WSIs for VessTrainSet, VessTestSet, and GlomTestSet 2

This human dataset was used to test adaptability of the model for vessels. In total there were 939 annotated arteries, 6023 arterioles, and 4507 glomeruli. *VessTrainSet* contained 226 renal tissue WSIs. *VessTestSet* contained an additional 58 holdout slides. Multiple stains per case were used. This dataset was manually annotated for relevant structures to establish a ground-truth.

The renal tissue WSIs came from UHC via co-author J.U.B. Diagnoses included thrombotic microangiopathy, hypertension-associated nephropathy, and vasculitis. Tissues were sectioned at 2-3 µm thickness. Diverse histologic stains were used, including H&E, PAS-H, Masson trichome, and Jones methenamine silver, for staining the tissue to depict different pathobiological features. A brightfield microscopy scanner Nanozoomer (Hamamatsu, Shizuoka, Japan) was used for WSI scanning at 40X resolution. Pixel resolution of the images used was 0.25 µm. Note that the *VessTestSet* dataset was used to construct the *GlomTestSet 2* dataset to conduct the study discussed in *GLOMERULI SEGMENTATION – scalability*.

### WSIs for IFTASet 1, IFTASet 2, IFTASet 3, IFTATestSet 2, and GlomTestSet 3

These datasets were used for the segmentation of IFTA. The human renal tissues for this part of the study came from four institutions: University of California, Davis; University of California, Los Angeles (UCLA); University of Coimbra (Portugal); and University Hospital Cologne (UHC).

Tissues were obtained from renal allograft nephropathy with no prior history of rejection. For this study, periodic acid-Schiff (PAS)-stained renal tissue WSIs of renal allograft nephropathy were used for training (*IFTASet 1, n=20; IFTASet 2, n=48;* and *IFTASet 3, n=22)*. One slide was selected per case for each institution. The WSIs per set were uniformly chosen from four IFTA classes defined based on semiquantitative score (ci/ct scores: 0, 1, 2, & 3); ci/ct scoring is a method defined in Banff 2018 criteria^18^ for assessing IFTA in transplant biopsies. A minimum of five slides per class were used for each set. The cases were reviewed to ensure the following selection criteria were met: (1) the amount of early or evolving IFTA with variable intermixed edema was minimized, (2) no active inflammation, (3) no prior history of rejection, and (4) cases were selected to represent the full range of IFTA severity. All types of IFTA, including classic, endocrinization, and thyroidization patterns, were included in the analysis, without distinguishing between the types. *IFTATestSet2* was provided by UHC, and contained 17 WSIs. This dataset followed similar case selection criteria as above with two slides from class 0 and five slides each from the remaining three classes.

The human data was collected in accordance with protocols approved by Institutional Review Boards at the UC Davis, UCLA, University of Coimbra, and the University at Buffalo. Deidentified images from UHC throughout this paper were used for retrospective research, and such is permitted under German law to conduct without IRB approval. The tissues were sectioned at 2-3 µm thickness and stained using PAS-H. Imaging was done using different brightfield microscopy WSI scanners, including Aperio CS virtual slide imaging system, Aperio AT2 (Leica Biosystems, Buffalo Grove, IL), and Nanozoomer (Hamamatsu, Shizuoka, Japan) at 40X resolution. Pixel resolution of the images used was 0.25 µm. Note that the *IFTATestSet 2* dataset was used to construct the *GlomTestSet 3* dataset to conduct the study discussed in *GLOMERULI SEGMENTATION – scalability*.

### KPMP WSI dataset

This dataset was used to test adaptability of the model for IFTA. This part of the study used 26 renal tissue biopsy whole slide images (WSIs) of 26 chronic kidney disease (CKD) subjects from Kidney Precision Medicine Project. The recruitment sites were Brigham & Women’s Hospital, Cleveland Clinic, Joslin Diabetes Center/ Beth Israel Deaconess Medical Center, and University of Texas at Southwestern. The inclusion criteria for CKD subjects for biopsy include subjects diagnosed with diabetic kidney disease (type 1 or 2) and hypertensive kidney disease. For the former, the subjects are included based on eGFR in the range of 30-59 mL/min/1.73 m^2^ or eGFR ≥ 60 with urinary protein to creatinine ratio (uPCR) > 150 mg/g or urinary albumin to creatinine ratio (uACR) > 30 mg/g. For the latter, the subjects are included based on eGFR in the range of 30-59 mL/min/1.73 m^2^ or eGFR ≥ 60 with uPCR in the range of 150-2000 mg/g or uACR in the range of 30-2000 mg/g. The study is overseen by three independent bodies, including a data safety monitoring board, a central institutional review board (WUSTL), and an NIH-NIDDK convened external expert panel. More details about the rationale and design of KPMP cases are available in a recent publication^44^. The tissues were sectioned at 2-3 µm thickness, and the PAS-H stained tissues were used for the study presented in this work. Imaging was done using Aperio GT450 brightfield microscopy WSI scanner (Leica Biosystems, Buffalo Grove, IL) at 40X resolution. Pixel resolution of the images used was 0.25 µm.

### WSIs for murine kidney tissue for the study discussed in MURINE MODEL ANALYSIS – utility

For this part of the study four murine model renal tissue WSIs were employed. These models include an aging model, two type 2 diabetic nephropathy (T2DN) models (KKAy and Db/Db), and a transgenic HIV-associated nephropathy (HIVAN) model. We used 8 mice (4 young and 4 old) WSIs for the aging model, 20 mice (10 KKAy or disease and 10 C57/BL6 or control) WSIs for the KKAy model, 14 mice (7 Db/Db or disease and 7 Db/m or wild-type control) WSIs for the Db/Db model, and 37 mice (26 transgenic disease and 11 wild-type control) WSIs for the HIVAN model.

The aging studies were performed in 4-month-old and 21-month-old C57/BL6 male mice obtained from the NIA aging rodent colony^45^. For the KKAy model (see published description^46^), male mice that develop spontaneous diabetes of polygenic origin were used. For the Db/Db model, male mice on BKS background featuring a leptin receptor mutation were used. These mice depict spontaneous/congenital diabetes due to leptin signaling abnormalities^47^. For the HIVAN model, transgenic Tg26 mice on a FVB/N background feature a *gag-pol*-deleted HIV-1 genome^48^ which manifests as collapsing glomerulopathy. Animal studies were performed in accordance with protocols approved by the Institutional Animal Care and Use Committee at the Georgetown University, National Institutes of Health, JHU, and UB, and are consistent with federal guidelines and regulations, and are in accordance with recommendations of the American Veterinary Medical Association guidelines on euthanasia. Tissues were sectioned at 2-3 µm thickness, and the PAS-H was used for staining. The slides were scanned using different brightfield microscopy WSI scanners, including Nanozoomer (Hamamatsu, Shizuoka, Japan) and MoticEasyScan Pro (Motic, San Antonio, TX), at 40X resolution. Pixel resolution of the images used was 0.25 µm.

### Software

With the goal of developing a tool with class leading WSI segmentation accuracy as well as easy accessibility to computational non-experts, we have integrated the popular semantic segmentation network Deeplab V3+^23^ with the DSA^13^, an open-source cloud-based histology management program. Specifically, we have created a suite of easy-to-use plugins using HistomicsUI, an application programing interface of the DSA for running Python codes. These plugins efficiently run the DeepLab network for native segmentation of WSIs, making testing new slides accessible through the HistomicsUI graphical user interface (the slide viewing component of the DSA). Using the HistomicsUI interface, users can interactively view the computational annotations, and further refine such annotations for training new models. The modified HistomicsTK-Deeplab codebase is available via GitHub and also as a prebuilt Docker image for easy installation. This software is deployed in the cloud and is accessible via the web, making it easily accessible to the community as a plug-and-play tool (***Fig. 1***). The open-source plugins are available to the digital pathology community for use and further development.

### Functionality

We have developed several plugin tools with various functions. (1) The *<SegmentWSI>* plugin (***Fig. 1a***) segments WSIs using a previously trained model. (2) The *<TrainNetwork>* plugin can be used to train new models from a folder of annotated WSIs (***Fig. 1b***). *Histo-Cloud* generates predictions as a series of image contours or sparse heatmaps which are written to JavaScript Object Notation (JSON) format for display in HistomicsUI as annotation layers. The code is modular, with the ability to handle multi-class segmentation, and includes the option to tweak the network hyperparameters for advanced users. (3) Functionality was included for conversion between JSON annotations and the XML format (*<IngestAperioXML>* and *<ExportAperioXML>* plugins). The XML format is used to display contours in Aperio ImageScope (Leica, Buffalo Grove, IL) which is a popular WSI viewer. (4) The *<ExtractFeaturesFromAnnotations>* plugin (see ***Fig. 1c***) was built for extraction of image and contour-based features from annotated regions in the slides. The features are written into the slide metadata (on DSA) in JSON format. For further data exploration, features saved into the slide metadata can be plotted pairwise using a scatterplot tool available in HistomicsUI (***Fig. 1c***) for a single slide or across a folder of WSIs. Features can also be saved in spreadsheet format for local download and further analysis.

### Computational model

We used the official implementation of the Deeplab V3+ segmentation network^23^, modified to work natively on WSIs. This implementation was accomplished by adapting the way the network ingests data, extracting patches from WSIs as needed during training using the *large_image* Python library^49^. A similar method (*HistoFetch*) is described more extensively in a recently published preprint^50^, which shows on-the-fly patch extraction speeds overall training time for unsupervised tasks. The *HistoFetch* method was adapted in this work to perform a supervised segmentation task by creating additional patch selection criteria intended to proactively balance uneven class distributions during patch extraction. Note that during development the code was migrated to use *large_image*^49^ for reading WSI data rather than the *openslide*^51^ library, as the former supports a larger number of slide formats. To convert the ground-truth annotations to masks for semantic segmentation, the HistomicsUI JSON annotations are converted into the Aperio ImageScope XML format, and the *XML_to_mask* conversion code from the original H-AI-L study^7^ was reused for generating ground-truth masks. This code follows the way *openslide* and *large_image* read WSI patches via specifying the location and scale of the patches. The min and max indices of each contour annotation are written into the metadata of the XML, allowing for faster reference of which contours are in the image region requested.

A flowchart providing an overview of this training input pipeline is presented in ***Supp. Fig. 1***. A similar pipeline is used during prediction (segmentation of slides), but patches are extracted deterministically from an overlapping grid pattern (excluding non-tissue regions) to ensure full tissue segmentation. The training and testing perform a fast color thresholding of the tissue region which is saved as a portable network graphics (PNG) mask for reference (to avoid repeated operations). This process ensures the network does not train on non-tissue regions, and thus speeds the prediction process. During the development, we found that occasionally providing the network with background (non-tissue) patches helped generalize the batch normalization parameters during training. We therefore implemented a parameter that defines the probability of selection of patches that may include the background region. A default of 0.1 was found to work well in generalizing the batch normalization layers.

### Training & Testing

Training of models was done on a server equipped with two Intel Xeon Silver 4114 (10 core) processors, with 64 GB RAM and dual Nvidia Quadro RTX 5000 graphical processing units (GPU) with 16 GB of video random access memory (VRAM). These resources allowed training with a batch size of 12 using image patches of size 512 × 512 pixels. A batch size of 12 is the minimum recommended for training the batch normalization parameters in the DeepLab implementation document. The *Athena* server (open for public use) has only one GPU with 8 GB of VRAM. We have therefore disabled training of the batch normalization parameters by default in the training plugin (which can be enabled in the advanced parameter section) and have set a default batch size of 2. All trained networks used a base learning rate of 1e^-3^ with polynomial decay using the momentum optimizer (momentum value=0.9).

All models use the *Xception 65* network backbone^23^, with DeepLab parameters *atrous_rates*=6, 12, and 18, *output_stride*=16, and *decoder_output_stride*=4 for both training and prediction. The *glomerulus model* was trained for 400,000 steps, and was initialized using the *ImageNet model*. The vessel segmentation models were trained for 100,000 steps, and the IFTA segmentation models were trained for 50,000 steps using *ImageNet model* as starting point for transfer learning. Details on the trained models are outlined in ***Table 1***.

As part of the input pipeline, WSI patches can be extracted efficiently at downsampled resolutions. The patch downsample rate is user specified, and multiple down sample rates can be specified during training, which are randomly cycled for patch extraction. For training, downsample rates of 1, 2, 3, & 4 with respect to the native slide resolution were used, a randomly selected downsample rate from the list was used for each extracted training patch. For prediction, a downsample rate of 2 was used for all experiments, we found this choice was a good compromise between prediction speed and accuracy. We believe that the multi-resolution training strategy helped the network to generalize. We found the *glomerulus model* works equally well in both 40X and 20X WSIs (both using a prediction downsample of 2). Further, the vessel segmentation model was trained using 40X WSIs, and successfully applied to the 20X GTEx WSIs for testing.

Using a large patch size for prediction increased segmentation performance, giving the network a larger field-of-view and reducing edge artifacts. For practical purposes, we settled on a default patch size of 2000 × 2000 pixels. For prediction, it was found that using a stride of 1000 pixels gave sufficient overlap between extracted patches. During prediction, the indices of the extracted patches are tracked, and the resulting bitmap prediction is used to populate a full WSI mask using the similar method as discussed in the original H-AI-L study^7^. To reduce the number of artifacts at the edge of the predicted patches, a parameter to remove the boarder of the predictions was included. Practically this parameter was set to remove 100 pixels from the border of each prediction.

To improve speed and to keep the memory requirements of code implementation low, network predictions are not up-sampled. Instead, the coordinates of the extracted contours or heatmap indices are up-sampled prior to JSON creation. Using DeepLab parameters, namely, *output_stride*=16 and *decoder_output_stride*=4, result in a prediction bitmap that is 25% of the size of the input resolution. With a default downsample of 2 used for prediction, the resultant WSI mask is one-eighth of the size of the pixel resolution of the original WSI. We found that 32 GB of RAM is enough to successfully segment even very large slides.

When experimenting with the network *logits* for the generation of the ROC plots (***Figs. 4a & 4b***), we converted the code to stitch the patch predictions together by averaging the *logits* of overlapping patches.

### Statistical analysis

Pearson correlation coefficient measure (Pearson’s *r*)^52^ was used for the study shown in ***Fig. 4c***, and corresponding *r* with null hypothesis *r=*0 vs alternative *r* > 0 was used to measure significance.

## AUTHOR CONTRIBUTIONS

B.L. wrote the *Histo-Cloud* code, created the plugins, performed the analysis, and wrote the manuscript. D.M. added features to HistomicsUI and offered advice for developing plugins. B.G. performed murine glomerular quantitation and assisted with glomeruli annotation. K.Y.J. organized the data for the IFTA study and annotated IFTA. L.R. and J.E.Z. provided data for the IFTA study and annotated IFTA. A.J.G. annotated IFTA. L.B. and C.E.A. conducted manual IFTA scoring in the KPMP biopsy cases. J.U.B. provided data for the IFTA and vessel segmentation tasks, and annotated arteries, arterioles, and IFTA. K.M. assisted J.U.B. in annotation. A.Z.R. organized the data for the murine model study, and also conducted manual IFTA scoring in the KPMP biopsy cases. X.X.W., K.M., B.A.J., M.L. generated KKAy, Db/Db, and aging cohorts. J.B.K. and T.Y. generated the HIVAN murine cohort, and J.B.K. edited the manuscript. S.S.H. and S.J. provided human renal tissue WSIs for training the *glomerulus model*. P.S. conceived the overall research plan, coordinated with the multi-disciplinary study team on the project and edited the manuscript.

## ACKNOWLEDGMENTS

This project was supported by NIH-NIDDK grant R01 DK114485 (PS), NIH-OD grants R01 DK114485 02S1 & 03S1 (PS), a pilot grant from the NIH-NIDDK CKD Biomarker Consortium grant U01 DK103225 (PS), via the opportunity pool funding mechanism, namely via glue grant mechanism (PS) of the NIH-NIDDK Kidney Precision Medicine Project grant U2C DK114886 (Contact: Dr. Jonathan Himmelfarb), a multi-disciplinary small team grant RSG201047.2 (PS) from the State University of New York, a pilot grant (PS) from the University of Buffalo’s Clinical and Translational Science Institute (CTSI) grant 3UL1TR00141206 S1 (Contact: Dr. Timothy Murphy), a DiaComp Pilot & Feasibility Project 21AU4180 (PS) with support from NIDDK Diabetic Complications Consortium grants U24 DK076169 and U24 DK115255 (Contact: Dr. Richard A. McIndoe), and NIH-OD’s HuBMAP grant U54 HL145608 (PS). The project was also supported by European Rare Kidney Disease Network and the Deutsche Forschungsgemeinschaft (BE-3801) (JUB), an intramural grant Cologne Fortune (KM), NIH-NIDDK R01 grants DK127830 & DK116567 (ML), and NIH-NIDDK Intramural Research Program (JBK). The KPMP CKD biopsy collection was supported by NIH-NIDDK grants UH3 DK114915 (Contact: Dr. Sushrut Waikar and Dr. Sylvia Rosas), UH3 DK114908 (Contact: Dr. Emilio Poggio), and UH3 DK114870 (Contact: Dr. Miguel Vazquez). We thank Seoul National University Hospital Human Biobank, a member of the National Biobank of Korea, which is supported by the Ministry of Health and Welfare, Republic of Korea. We thank Dr. Agnes Fogo for providing the human renal biopsy WSIs from VUMC which were used in our earlier publications and are reused in this work for training the glomerulus model. We thank Dr. Rabi Yacoub for generating the STZ and nephrin KD murine models which were used in our earlier publications and are reused in this work for training the glomerulus model. We thank Ms. Briana Santo, Ms. Darshana Govind, Mr. Nicholas Lucarelli, and Mr. Samuel Boarder (graduate students of Dr. Sarder) for their help in glomeruli annotations for training the glomerulus model. We also thank Ms. Stephanie Grewenow and Ms. Becky Steck for their help in organizing the KPMP renal tissue biopsy WSIs.v

## SUPPLEMENTAL MATERIAL

Supplemental Figure 1. Flowchart of the custom DeepLab WSI input pipeline.

Supplemental Figure 2. Glomeruli segmentation performance using IFTA segmentation models.

Supplemental Figure 3. ROC performance for glomeruli and vessel segmentation.

Supplemental Figure 4. Correlation of percent IFTA estimation between methods.

Supplemental Figure 5. Quantifying UMAP separability for the murine model studies.

Supplemental Table 1. Table of abbreviations.

Supplemental Table 2. Features measured on each glomerulus.

