## Supplemental Material for "A user-friendly tool for cloud-based whole slide image segmentation, with examples from renal histopathology"

### SUPPLIMENTAL FIGURES

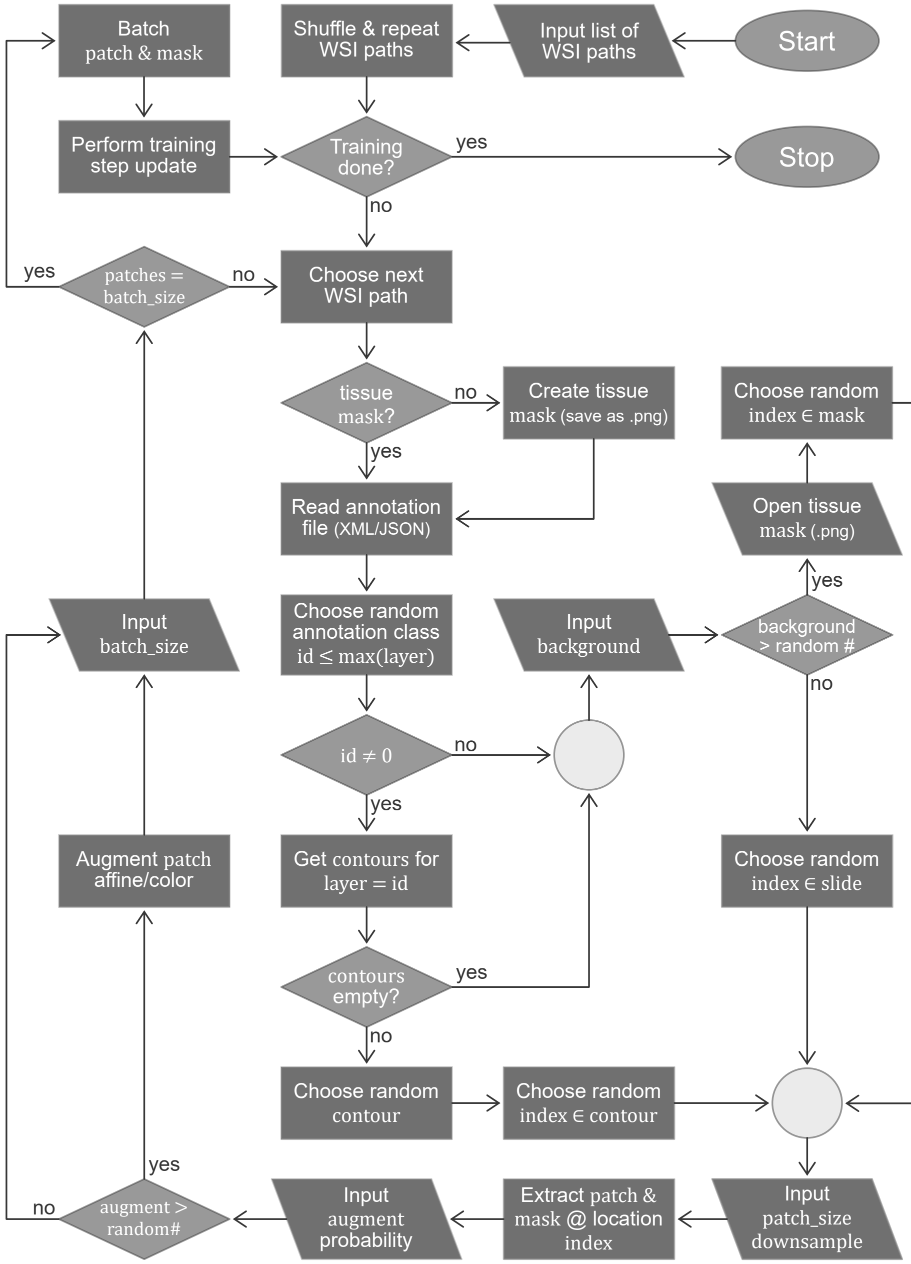


Supplemental Fig. 1 | Flowchart of the custom DeepLab WSI input pipeline.

The details of the custom input pipeline used by our modified DeepLab code to ingest WSI data during training. The *large_image* python library (https://github.com/girder/large_image/) is used to extract patches from WSIs on-the-fly. This process uses a modified version of our *HistoFetch* pipeline[1], which has been further modified to work for supervised learning tasks. For the network training, pixel locations from the image data corresponding to each data class are randomly selected by exploiting the XML or JSON annotation files. This ensures class balancing for network training by selecting appropriately sampled pixel regions for all the classes. If the background class is selected, a random location within the tissue region (which has been pre-segmented via morphological processing) is selected. During application development, we found that occasionally providing the network with non-tissue patches as background helped the batch normalization parameters to generalize, which reduced error. We therefore added a parameter defining the probability of selection of a non-tissue region, allowing patches within and outside the tissue regions to be included in the analyses. When using a trained model to segment structures (prediction on new slides), a similar pipeline is used. However, image patches to be processed are extracted deterministically from an overlapping grid pattern (excluding non-tissue regions), to ensure the entire tissue region is processed for full segmentation. This input pipeline is predominantly implemented in the following files in the DeepLab codebase available via github: /datasets/wsi_data_generator.py and /utils/wsi_dataset_util_large_image.py

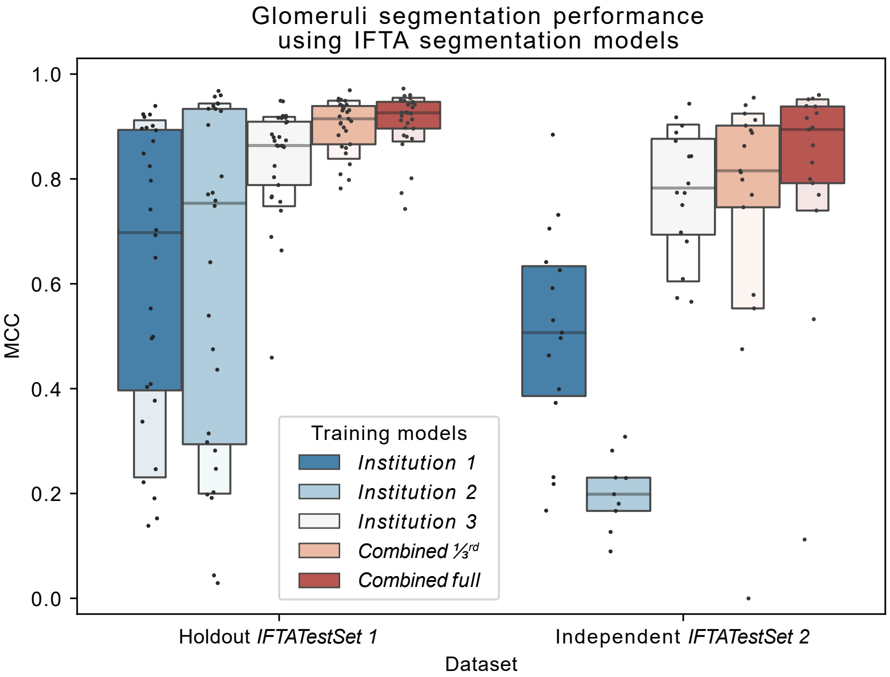


***Supplemental Fig. 2* | Glomeruli segmentation performance using IFTA segmentation models.**
The glomerular segmentation performance using the five models trained for segmenting IFTA and glomeruli (see *IFTA SEGMENTATION – adaptability* under *RESULTS* and ***Fig. 4***). The performance is quantified on the *IFTATestSet 1* with 29 holdout renal tissue WSIs, and the independent test set *IFTATestSet 2* with 17 renal tissue WSIs and annotation ground-truth originated from an institution independent of the training dataset and *IFTATestSet 1*. We observe the same trend in performance as IFTA segmentation as shown in ***Fig. 4***. Namely, the *Combined full* model delivers the best performance, while the *Combined 1/3rd* model performs better than any of the models trained on a single institution data alone.

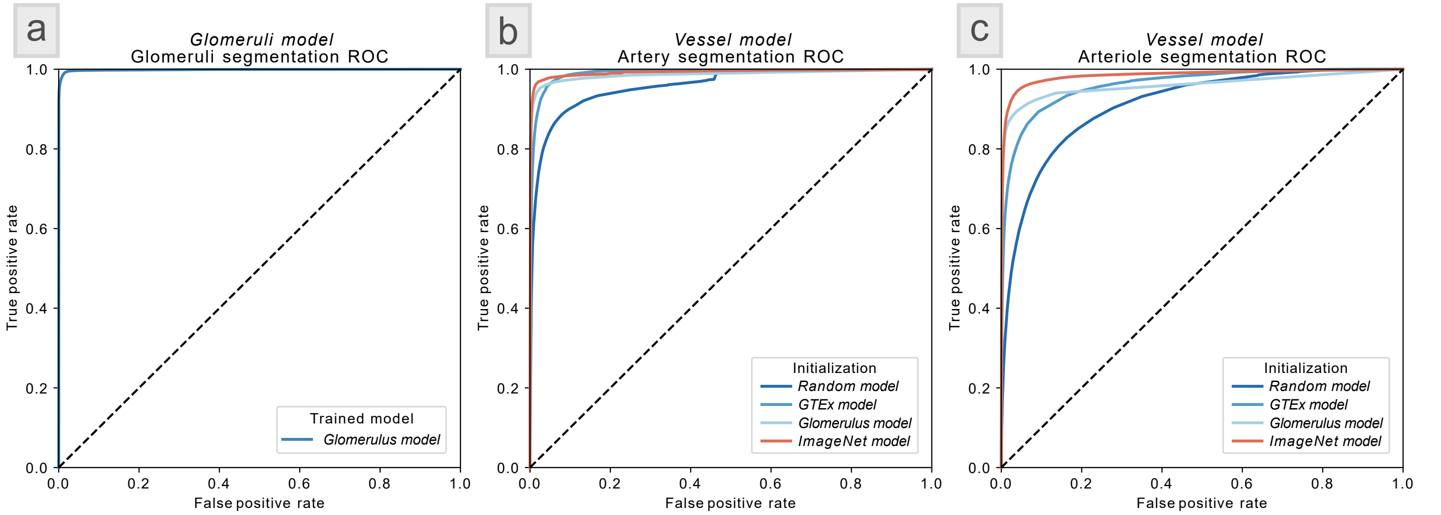


***Supplemental Fig. 3* | ROC performance for glomeruli and vessel segmentation.**

**Panel [a]** shows the ROC performance for glomerular segmentation on *GlomTestSet 1* with 100 holdout renal tissue WSIs using *glomerular model* (see *GLOMERULI SEGMENTATION – scalability* in *RESULTS* and ***Fig. 2***). **[b]** shows the ROC performance for artery segmentation on the holdout dataset *VessTestSet* with 58 renal tissue WSIs using the four initialization strategies for vessel segmentation (see *VESSEL SEGMENTATION – adaptability* in *RESULTS* and ***Fig. 3***). **[c]** shows the ROC performance for arteriole segmentation on the same holdout dataset with same initialization strategies as used in **[b]**.

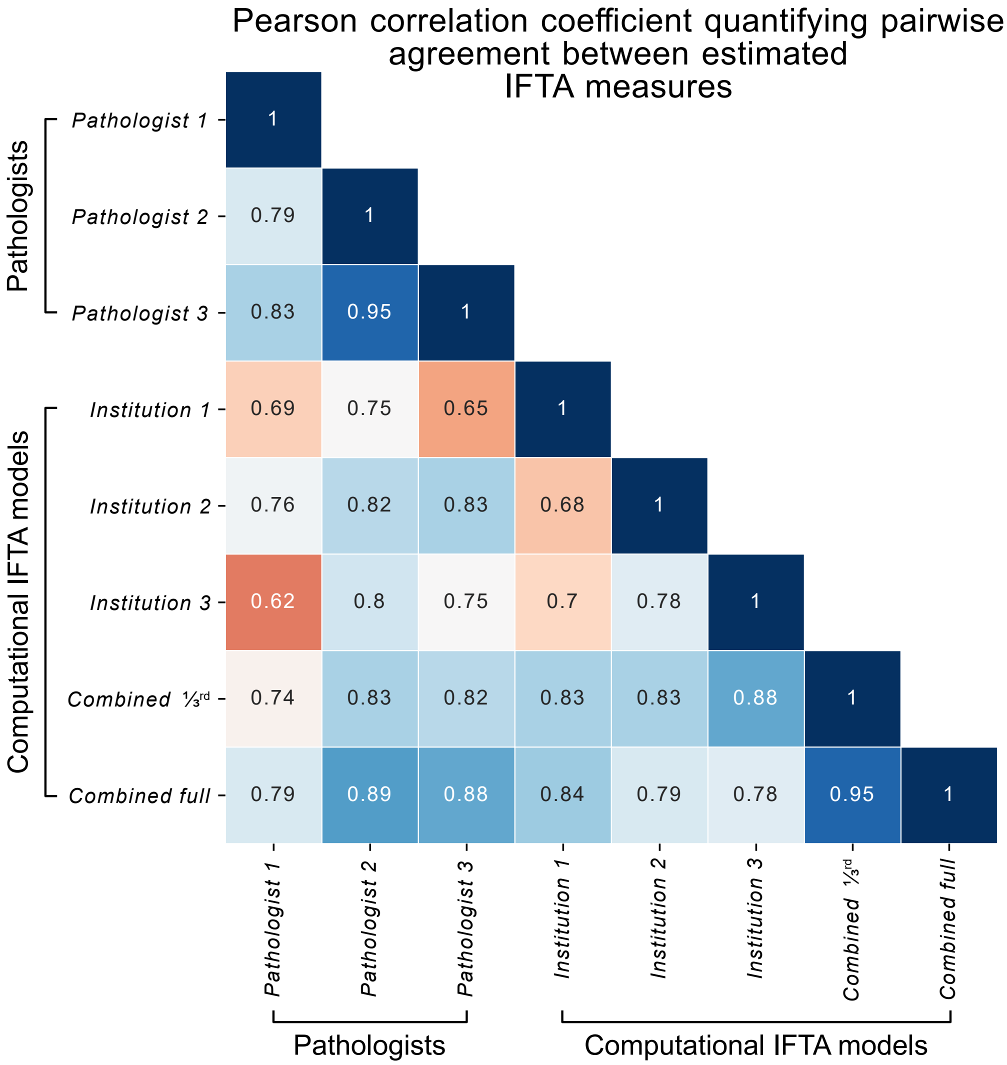


***Supplemental Fig. 4* | Correlation of percent IFTA estimation between methods.**

Pearson correlation coefficient quantifying pairwise agreement between computationally and manually estimated IFTA measures for 26 CKD renal tissue WSIs from Kidney Precision Medicine Project (KPMP) cohort data, *KPMPTestSet*. The full confusion matrix of correlations of computationally estimated and visually manually estimated percent IFTA for the KPMP test cohort. Pairwise correlations were measured for each pair of the three KPMP pathologists and the five computational models trained for IFTA segmentation (see *IFTA SEGMENTATION – adaptability* under *RESULTS*). This result is an extension of ***Fig. 4c***. Pearson correlation coefficients with $p$-value < 0.05 are shown.

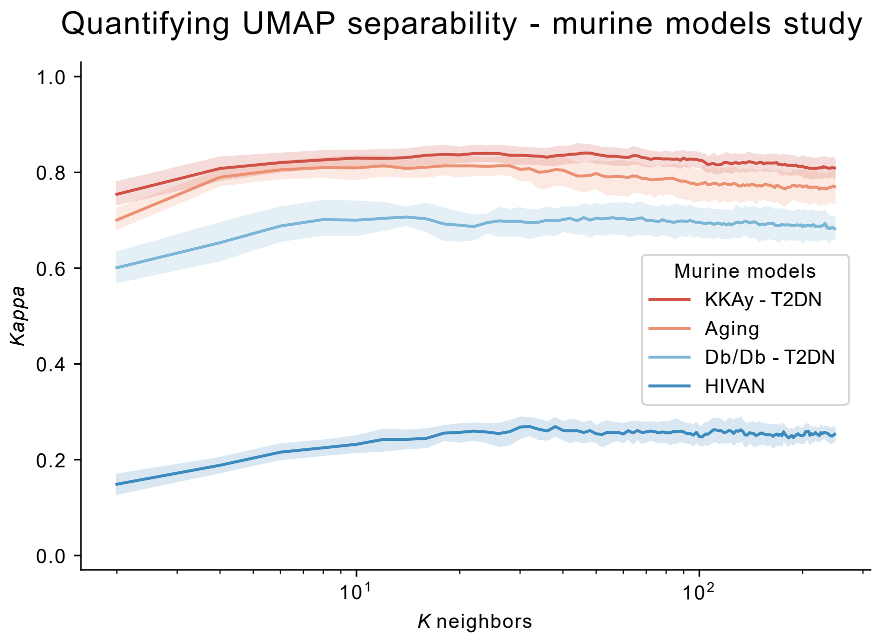


***Supplemental Fig. 5*** **|** Quantifying UMAP separability for the murine model studies.
K-nearest neighbors (KNN) classifier performance plotting the *Cohen's Kappa* measure as a function of *K* neighbors for classifying the unsupervised UMAP features with respect to disease vs control status for the murine kidney disease models (see ***Fig. 5***). This analysis was done using 10-fold cross validation using a similar method as formalized in a previous work[2].

### TABLE OF ABBREVIATIONS

Supplementary Table 1 | Features measured on each glomerulus.

| | |
|---|---|
| AUC | Area under the curve |
| CKD | Chronic kidney disease |
| CNN | Convolutional neural network |
| DSA | Digital slide archive |
| eGFR | Estimated glomerular filtration rate |
| FSGS | Focal segmental glomerulosclerosis |
| GPU | G**raphics processing unit** |
| GTEx | Genotype-tissue expression |
| H-AI-L | Human – Artificial Intelligence – Loop |
| H&E | Hematoxylin and eosin (stain) |
| HIVAN | HIV-associated nephropathy |
| IFTA | Interstitial fibrosis and tubular atrophy |
| IoU | Intersection over Union (Jaccard index) |
| JSON | JavaScript object notation |
| KD | Knockdown |
| KPMP | Kidney precision medicine project |
| MCC | Matthews correlation coefficient |
| PAS | Periodic acid–Schiff (stain) |
| PNG | Portable network graphics |
| RAM | R**andom access memory** |
| ROC | Receiver operating characteristic |
| STZ | Streptozotocin |
| T2DN | Type 2 diabetic nephropathy |
| uACR | Albumin to creatinine ratio |
| UMAP | Uniform manifold approximation and projection |
| uPCR | Urinary protein to creatinine ratio |
| VRAM | V**ideo RAM** |
| WSI | Whole slide image |
| XML | Extensible markup language |

### GLOMERULAR FEATURES

#### Supplementary Table 2 | Features measured on each glomerulus.

Note that each segmented glomerulus is further computationally sub-compartmentalized for PAS+ area, nuclei, and luminal white spaces, and the features are subsequently quantified from these sub-compartments using a method as discussed in our previous work3. In the list below C represents the color features, D the distance features, M the morphological features, T the textural features, and PAS+ the periodic acid-Schiff positive features.

| Index | Feature Name | Type |
|---|---|---|
| 1 | Mean of red values in PAS+ regions | C |
| 2 | Mean of green values in PAS+ regions | C |
| 3 | Mean of blue values in PAS+ regions | C |
| 4 | Standard deviation of red values in PAS+ regions | C |
| 5 | Standard deviation of green values in PAS+ regions | C |
| 6 | Standard deviation of blue values in PAS+ regions | C |
| 7 | Mean of red values in luminal regions | C |
| 8 | Mean of green values in luminal regions | C |
| 9 | Mean of blue values in luminal regions | C |
| 10 | Standard deviation of red values in luminal regions | C |
| 11 | Standard deviation of green values in luminal regions | C |
| 12 | Standard deviation of blue values in luminal regions | C |
| 13 | Mean of red values in nuclear regions | C |
| 14 | Mean of green values in nuclear regions | C |
| 15 | Mean of blue values in nuclear regions | C |
| 16 | Standard deviation of red values in nuclear regions | C |
| 17 | Standard deviation of green values in nuclear regions | C |
| 18 | Standard deviation of blue values in nuclear regions | C |
| 19 | Mean distance of luminal object centroids from glomerular centroid | D |
| 20 | Mean of mean distances of luminal object centroids from glomerular boundary | D |
| 21 | Mean of maximum distances of luminal object centroids from glomerular boundary | D |
| 22 | Mean of minimum distances of luminal object centroids from glomerular boundary | D |
| 23 | Mean of mean distances of luminal object centroids from themselves | D |
| 24 | Mean of maximum distances of luminal object centroids from themselves | D |
| 25 | Mean of minimum distances of luminal object centroids from themselves | D |
| 26 | Mean distance of PAS+ object centroids from glomerular centroid | D |
| 27 | Mean of mean distances of PAS+ object centroids from glomerular boundary | D |
| 28 | Mean of maximum distances of PAS+ object centroids from glomerular boundary | D |
| 29 | Mean of minimum distances of PAS+ object centroids from glomerular boundary | D |
| 30 | Mean of mean distances of PAS+ object centroids from themselves | D |
| 31 | Mean of maximum distances of PAS+ object centroids from themselves | D |
| 32 | Mean of minimum distances of PAS+ object centroids from themselves | D |
| 33 | Mean distance of nuclear object centroids from glomerular centroid | D |
| 34 | Mean of mean distances of nuclear object centroids from glomerular boundary | D |
| 35 | Mean of maximum distances of nuclear object centroids from glomerular boundary | D |
| 36 | Mean of minimum distances of nuclear object centroids from glomerular boundary | D |
| 37 | Mean of mean distances of nuclear object centroids from themselves | D |
| 38 | Mean of maximum distances of nuclear object centroids from themselves | D |
| 39 | Mean of minimum distances of nuclear object centroids from themselves | D |
| 40 | Average luminal object solidity | M |
| 41 | Average PAS+ region contained in luminal object boundaries | M |

| | | |
|---|---|---|
| 42 | Average nuclear region contained in luminal object boundaries | M |
| 43 | Sum total luminal objects' areas | M |
| 44 | Mean of luminal objects' areas | M |
| 45 | Median of luminal objects' areas | M |
| 46 | Average PAS+ object solidity | M |
| 47 | Average lumina region contained in PAS+ object boundaries | M |
| 48 | Average nuclear region contained in PAS+ object boundaries | M |
| 49 | Sum total PAS+ objects' areas | M |
| 50 | Mean of PAS+ objects' areas | M |
| 51 | Median of PAS+ objects' areas | M |
| 52 | Mean ratio of PAS+ pixels lying just outside nuclear perimeter to length of perimeter | M |
| 53 | Mean ratio of luminal pixels lying just outside nuclear perimeter to length of perimeter | M |
| 54 | Mean nuclear perimeter pixel count | M |
| 55 | Sum total nuclear area | M |
| 56 | Mean nuclear areas | M |
| 57 | Mode nuclear areas | M |
| 58 | Total glomerular area | M |
| 59 | Total PAS+ object number | M |
| 60 | Total luminal object number | M |
| 61 | Total nucleus number | M |
| 62 | Sum of PAS+ distance transform values 0µm < d ≤ 2.5µm | M |
| 63 | Sum of PAS+ distance transform values 2.5µm < d ≤ 5µm | M |
| 64 | Sum of PAS+ distance transform values 5µm < d ≤ 250µm | M |
| 65 | Maximum PAS+ distance transform value 2.5µm < d ≤ 5µm | M |
| 66 | Number of connected objects with PAS+ distance transform value 0µm < d ≤ 2.5µm | M |
| 67 | Number of connected objects with PAS+ distance transform value 2.5 µm < d ≤ 5µm | M |
| 68 | Mean of PAS+ distance transform values 0µm < d ≤ 2.5µm | M |
| 69 | Mean of PAS+ distance transform values 2.5µm < d ≤ 5µm | M |
| 70 | Median of PAS+ distance transform values 0µm < d ≤ 2.5µm | M |
| 71 | Median of PAS+ distance transform values 2.5µm < d ≤ 5µm | M |
| 72 | Mean area of objects with PAS+ distance transform value 0µm < d ≤ 2.5µm | M |
| 73 | Median area of objects with PAS+ distance transform value 0µm < d ≤ 2.5µm | M |
| 74 | Maximum area of objects with PAS+ distance transform value 0µm < d ≤ 2.5µm | M |
| 75 | Mean area of objects with PAS+ distance transform value 2.5µm < d ≤ 5µm | M |
| 76 | Median area of objects with PAS+ distance transform value 2.5µm < d ≤ 5µm | M |
| 77 | Count of pixels with PAS+ distance transform value 0.25µm < d ≤ 0.75µm | M |
| 78 | Count of pixels with PAS+ distance transform value 0.75µm < d ≤ 1.25µm | M |
| 79 | Count of pixels with PAS+ distance transform value 1.25µm < d ≤ 1.75µm | M |
| 80 | Count of pixels with PAS+ distance transform value 1.75µm < d ≤ 2.25µm | M |
| 81 | Count of pixels with PAS+ distance transform value 2.25µm < d ≤ 2.75µm | M |
| 82 | Count of pixels with PAS+ distance transform value 2.75µm < d ≤ 3.25µm | M |
| 83 | Count of pixels with PAS+ distance transform value 3.25µm < d ≤ 3.75µm | M |
| 84 | Count of pixels with PAS+ distance transform value 3.75µm < d ≤ 4.25µm | M |
| 85 | Count of pixels with PAS+ distance transform value 4.25µm < d ≤ 4.75µm | M |
| 86 | Count of pixels with PAS+ distance transform value 4.75µm < d ≤ 5.25µm | M |
| 87 | Count of pixels with PAS+ distance transform value 5.25µm < d ≤ 5.75µm | M |
| 88 | Count of pixels with PAS+ distance transform value 5.75µm < d ≤ 6.25µm | M |

| | | |
|---|---|---|
| 89 | Count of pixels with PAS+ distance transform value 6.25µm < d ≤ 6.75µm | M |
| 90 | Count of pixels with PAS+ distance transform value 6.75µm < d ≤ 7.25µm | M |
| 91 | Count of pixels with PAS+ distance transform value 7.25µm < d ≤ 7.75µm | M |
| 92 | Count of pixels with PAS+ distance transform value 7.75µm < d ≤ 8.25µm | M |
| 93 | Count of pixels with PAS+ distance transform value 8.25µm < d ≤ 8.75µm | M |
| 94 | Count of pixels with PAS+ distance transform value 8.75µm < d ≤ 9.25µm | M |
| 95 | Count of pixels with PAS+ distance transform value 9.25µm < d ≤ 9.75µm | M |
| 96 | Count of pixels with PAS+ distance transform value 9.75µm < d ≤ 10.25µm | M |
| 97 | Count of pixels with PAS+ distance transform value 10.25µm < d ≤ 10.75µm | M |
| 98 | Count of pixels with PAS+ distance transform value 10.75µm < d ≤ 11.25µm | M |
| 99 | Count of pixels with PAS+ distance transform value 11.25µm < d ≤ 11.75µm | M |
| 100 | Count of pixels with PAS+ distance transform value 11.75µm < d ≤ 12.25µm | M |
| 101 | Count of pixels with PAS+ distance transform value 12.25µm < d ≤ 12.75µm | M |
| 102 | Count of pixels with PAS+ distance transform value 12.75µm < d ≤ 13.25µm | M |
| 103 | Count of pixels with PAS+ distance transform value 13.25µm < d ≤ 13.75µm | M |
| 104 | Count of pixels with PAS+ distance transform value 13.75µm < d ≤ 14.25µm | M |
| 105 | Count of pixels with PAS+ distance transform value 14.25µm < d ≤ 14.75µm | M |
| 106 | Count of pixels with PAS+ distance transform value 14.75µm < d ≤ 15.25µm | M |
| 107 | Count of pixels with PAS+ distance transform value 15.25µm < d ≤ 15.75µm | M |
| 108 | Count of pixels with PAS+ distance transform value 15.75µm < d ≤ 16.25µm | M |
| 109 | Count of pixels with PAS+ distance transform value 16.25µm < d ≤ 16.75µm | M |
| 110 | Count of pixels with PAS+ distance transform value 16.75µm < d ≤ 17.25µm | M |
| 111 | Count of pixels with PAS+ distance transform value 17.25µm < d ≤ 17.75µm | M |
| 112 | Count of pixels with PAS+ distance transform value 17.75µm < d ≤ 18.25µm | M |
| 113 | Count of pixels with PAS+ distance transform value 18.25µm < d ≤ 18.75µm | M |
| 114 | Count of pixels with PAS+ distance transform value 18.75µm < d ≤ 19.25µm | M |
| 115 | Count of pixels with PAS+ distance transform value 19.25µm < d ≤ 19.75µm | M |
| 116 | Count of pixels with PAS+ distance transform value 19.75µm < d ≤ 500µm | M |
| 117 | Count of pixels with luminal distance transform value 0.25µm < d ≤ 0.5µm | M |
| 118 | Count of pixels with luminal distance transform value 0.5µm < d ≤ 0.75µm | M |
| 119 | Count of pixels with luminal distance transform value 0.75µm < d ≤ 1µm | M |
| 120 | Count of pixels with luminal distance transform value 1µm < d ≤ 1.25µm | M |
| 121 | Count of pixels with luminal distance transform value 1.25µm < d ≤ 1.5µm | M |
| 122 | Count of pixels with luminal distance transform value 1.5µm < d ≤ 1.75µm | M |
| 123 | Count of pixels with luminal distance transform value 1.75µm < d ≤ 2µm | M |
| 124 | Count of pixels with luminal distance transform value 2µm < d ≤ 2.25µm | M |
| 125 | Count of pixels with luminal distance transform value 2.25µm < d ≤ 2.5µm | M |
| 126 | Count of pixels with luminal distance transform value 2.5µm < d ≤ 2.75µm | M |
| 127 | Count of pixels with luminal distance transform value 2.75µm < d ≤ 3µm | M |
| 128 | Count of pixels with luminal distance transform value 3µm < d ≤ 3.25µm | M |
| 129 | Count of pixels with luminal distance transform value 3.25µm < d ≤ 3.5µm | M |
| 130 | Count of pixels with luminal distance transform value 3.5µm < d ≤ 3.75µm | M |
| 131 | Count of pixels with luminal distance transform value 3.75µm < d ≤ 4µm | M |
| 132 | Count of pixels with luminal distance transform value 4µm < d ≤ 4.25µm | M |
| 133 | Count of pixels with luminal distance transform value 4.25µm < d ≤ 4.5µm | M |
| 134 | Count of pixels with luminal distance transform value 4.5µm < d ≤ 4.75µm | M |
| 135 | Count of pixels with luminal distance transform value 4.75µm < d ≤ 5µm | M |
| 136 | Count of pixels with luminal distance transform value 5µm < d ≤ 5.25µm | M |

| | | |
|---|---|---|
| 137 | Count of pixels with luminal distance transform value 5.25µm < d ≤ 5.5µm | M |
| 138 | Count of pixels with luminal distance transform value 5.5µm < d ≤ 5.75µm | M |
| 139 | Count of pixels with luminal distance transform value 5.75µm < d ≤ 6µm | M |
| 140 | Count of pixels with luminal distance transform value 6µm < d ≤ 6.25µm | M |
| 141 | Count of pixels with luminal distance transform value 6.25µm < d ≤ 6.5µm | M |
| 142 | Count of pixels with luminal distance transform value 6.5µm < d ≤ 6.75µm | M |
| 143 | Count of pixels with luminal distance transform value 6.75µm < d ≤ 7µm | M |
| 144 | Count of pixels with luminal distance transform value 7µm < d ≤ 7.25µm | M |
| 145 | Count of pixels with luminal distance transform value 7.25µm < d ≤ 7.5µm | M |
| 146 | Count of pixels with luminal distance transform value 7.5µm < d ≤ 7.75µm | M |
| 147 | Count of pixels with luminal distance transform value 7.75µm < d ≤ 8µm | M |
| 148 | Count of pixels with luminal distance transform value 8µm < d ≤ 8.25µm | M |
| 149 | Count of pixels with luminal distance transform value 8.25µm < d ≤ 8.5µm | M |
| 150 | Count of pixels with luminal distance transform value 8.5µm < d ≤ 8.75µm | M |
| 151 | Count of pixels with luminal distance transform value 8.75µm < d ≤ 9µm | M |
| 152 | Count of pixels with luminal distance transform value 9µm < d ≤ 9.25µm | M |
| 153 | Count of pixels with luminal distance transform value 9.25µm < d ≤ 9.5µm | M |
| 154 | Count of pixels with luminal distance transform value 9.5µm < d ≤ 9.75µm | M |
| 155 | Count of pixels with luminal distance transform value 9.75µm < d ≤ 10µm | M |
| 156 | Count of pixels with luminal distance transform value 10µm < d ≤ 10.25µm | M |
| 157 | Count of pixels with luminal distance transform value 10.25µm < d ≤ 10.5µm | M |
| 158 | Count of pixels with luminal distance transform value 10.5µm < d ≤ 10.75µm | M |
| 159 | Count of pixels with luminal distance transform value 10.75µm < d ≤ 11µm | M |
| 160 | Count of pixels with luminal distance transform value 11µm < d ≤ 11.25µm | M |
| 161 | Count of pixels with luminal distance transform value 11.25µm < d ≤ 11.5µm | M |
| 162 | Count of pixels with luminal distance transform value 11.5µm < d ≤ 11.75µm | M |
| 163 | Count of pixels with luminal distance transform value 11.75µm < d ≤ 12µm | M |
| 164 | Count of pixels with luminal distance transform value 12µm < d ≤ 12.25µm | M |
| 165 | Count of pixels with luminal distance transform value 12.25µm < d ≤ 12.5µm | M |
| 166 | Count of pixels with luminal distance transform value 12.5µm < d ≤ 12.75µm | M |
| 167 | Count of pixels with luminal distance transform value 12.75µm < d ≤ 13µm | M |
| 168 | Count of pixels with luminal distance transform value 13µm < d ≤ 13.25µm | M |
| 169 | Count of pixels with luminal distance transform value 13.25µm < d ≤ 13.5µm | M |
| 170 | Count of pixels with luminal distance transform value 13.5µm < d ≤ 13.75µm | M |
| 171 | Count of pixels with luminal distance transform value 13.75µm < d ≤ 14µm | M |
| 172 | Count of pixels with luminal distance transform value 14µm < d ≤ 14.25µm | M |
| 173 | Count of pixels with luminal distance transform value 14.25µm < d ≤ 14.5µm | M |
| 174 | Count of pixels with luminal distance transform value 14.5µm < d ≤ 14.75µm | M |
| 175 | Count of pixels with luminal distance transform value 14.75µm < d ≤ 15µm | M |
| 176 | Count of pixels with luminal distance transform value 15µm < d ≤ 500µm | M |
| 177 | Count of pixels with nuclear distance transform value 0.25µm < d ≤ 0.5µm | M |
| 178 | Count of pixels with nuclear distance transform value 0.5µm < d ≤ 0.75µm | M |
| 179 | Count of pixels with nuclear distance transform value 0.75µm < d ≤ 1µm | M |
| 180 | Count of pixels with nuclear distance transform value 1µm < d ≤ 1.25µm | M |
| 181 | Count of pixels with nuclear distance transform value 1.25µm < d ≤ 1.5µm | M |
| 182 | Count of pixels with nuclear distance transform value 1.5µm < d ≤ 1.75µm | M |
| 183 | Count of pixels with nuclear distance transform value 1.75µm < d ≤ 2µm | M |
| 184 | Count of pixels with nuclear distance transform value 2µm < d ≤ 2.25µm | M |

| | | |
|---|---|---|
| 185 | Count of pixels with nuclear distance transform value 2.25µm < d ≤ 2.5µm | M |
| 186 | Count of pixels with nuclear distance transform value 2.5µm < d ≤ 2.75µm | M |
| 187 | Count of pixels with nuclear distance transform value 2.75µm < d ≤ 3µm | M |
| 188 | Count of pixels with nuclear distance transform value 3µm < d ≤ 3.25µm | M |
| 189 | Count of pixels with nuclear distance transform value 3.25µm < d ≤ 3.5µm | M |
| 190 | Count of pixels with nuclear distance transform value 3.5µm < d ≤ 3.75µm | M |
| 191 | Count of pixels with nuclear distance transform value 3.75µm < d ≤ 4µm | M |
| 192 | Count of pixels with nuclear distance transform value 4µm < d ≤ 4.25µm | M |
| 193 | Count of pixels with nuclear distance transform value 4.25µm < d ≤ 4.5µm | M |
| 194 | Count of pixels with nuclear distance transform value 4.5µm < d ≤ 4.75µm | M |
| 195 | Count of pixels with nuclear distance transform value 4.75µm < d ≤ 5µm | M |
| 196 | Count of pixels with nuclear distance transform value 5µm < d ≤ 500µm | M |
| 197 | Count of pixels with glomerular distance transform value 0.5µm < d ≤ 6.75µm | M |
| 198 | Count of pixels with glomerular distance transform value 6.75µm < d ≤ 13µm | M |
| 199 | Count of pixels with glomerular distance transform value 13µm < d ≤ 19.25µm | M |
| 200 | Count of pixels with glomerular distance transform value 19.25µm < d ≤ 25.5µm | M |
| 201 | Count of pixels with glomerular distance transform value 25.5µm < d ≤ 31.75µm | M |
| 202 | Count of pixels with glomerular distance transform value 31.75µm < d ≤ 38µm | M |
| 203 | Count of pixels with glomerular distance transform value 38µm < d ≤ 44.25µm | M |
| 204 | Count of pixels with glomerular distance transform value 44.25µm < d ≤ 50.5µm | M |
| 205 | Count of pixels with glomerular distance transform value 50.5µm < d ≤ 56.75µm | M |
| 206 | Count of pixels with glomerular distance transform value 56.75µm < d ≤ 63µm | M |
| 207 | Count of pixels with glomerular distance transform value 63µm < d ≤ 69.25µm | M |
| 208 | Count of pixels with glomerular distance transform value 69.25µm < d ≤ 75.5µm | M |
| 209 | Count of pixels with glomerular distance transform value 75.5µm < d ≤ 81.75µm | M |
| 210 | Count of pixels with glomerular distance transform value 81.75µm < d ≤ 88µm | M |
| 211 | Count of pixels with glomerular distance transform value 88µm < d ≤ 94.25µm | M |
| 212 | Count of pixels with glomerular distance transform value 94.25µm < d ≤ 100.5µm | M |
| 213 | Count of pixels with glomerular distance transform value 100.5µm < d ≤ 106.75µm | M |
| 214 | Count of pixels with glomerular distance transform value 106.75µm < d ≤ 113µm | M |
| 215 | Count of pixels with glomerular distance transform value 113µm < d ≤ 119.25µm | M |
| 216 | Count of pixels with glomerular distance transform value 119.25µm < d ≤ 125.5µm | M |
| 217 | Count of pixels with glomerular distance transform value 125.5µm < d ≤ 131.75µm | M |
| 218 | Count of pixels with glomerular distance transform value 131.75µm < d ≤ 138µm | M |
| 219 | Count of pixels with glomerular distance transform value 138µm < d ≤ 144.25µm | M |
| 220 | Count of pixels with glomerular distance transform value 144.25µm < d ≤ 5000µm | M |
| 221 | Number nuclear pixels contained radius 0µm < R ≤ 25µm | M |
| 222 | Number nuclear pixels contained radius 25µm < R ≤ 50µm | M |
| 223 | Number nuclear pixels contained radius 50µm < R ≤ 75µm | M |
| 224 | Number nuclear pixels contained radius 75µm < R ≤ 100µm | M |
| 225 | Number nuclear pixels contained radius 100µm < R ≤ 125µm | M |
| 226 | Number nuclear pixels contained radius 125µm < R ≤ 150µm | M |
| 227 | Number nuclear pixels contained radius 150µm < R ≤ 175µm | M |
| 228 | Number nuclear pixels contained radius 175µm < R ≤ 200µm | M |
| 229 | Number nuclear pixels contained radius 200µm < R ≤ 225µm | M |
| 230 | Number nuclear pixels contained radius 225µm < R ≤ 250µm | M |
| 231 | Number nuclear pixels contained radius 250µm < R ≤ 325µm | M |
| 232 | Number luminal pixels contained radius 0µm < R ≤ 25µm | M |

| | | |
|---|---|---|
| 233 | Number luminal pixels contained radius 25µm < R ≤ 50µm | M |
| 234 | Number luminal pixels contained radius 50µm < R ≤ 75µm | M |
| 235 | Number luminal pixels contained radius 75µm < R ≤ 100µm | M |
| 236 | Number luminal pixels contained radius 100µm < R ≤ 125µm | M |
| 237 | Number luminal pixels contained radius 125µm < R ≤ 150µm | M |
| 238 | Number luminal pixels contained radius 150µm < R ≤ 175µm | M |
| 239 | Number luminal pixels contained radius 175µm < R ≤ 200µm | M |
| 240 | Number luminal pixels contained radius 200µm < R ≤ 225µm | M |
| 241 | Number luminal pixels contained radius 225µm < R ≤ 250µm | M |
| 242 | Number luminal pixels contained radius 250µm < R ≤ 325µm | M |
| 243 | Number PAS+ pixels contained radius 0µm < R ≤ 25µm | M |
| 244 | Number PAS+ pixels contained radius 25µm < R ≤ 50µm | M |
| 245 | Number PAS+ pixels contained radius 50µm < R ≤ 75µm | M |
| 246 | Number PAS+ pixels contained radius 75µm < R ≤ 100µm | M |
| 247 | Number PAS+ pixels contained radius 100µm < R ≤ 125µm | M |
| 248 | Number PAS+ pixels contained radius 125µm < R ≤ 150µm | M |
| 249 | Number PAS+ pixels contained radius 150µm < R ≤ 175µm | M |
| 250 | Number PAS+ pixels contained radius 175µm < R ≤ 200µm | M |
| 251 | Number PAS+ pixels contained radius 200µm < R ≤ 225µm | M |
| 252 | Number PAS+ pixels contained radius 225µm < R ≤ 250µm | M |
| 253 | Number PAS+ pixels contained radius 250µm < R ≤ 325µm | M |
| 254 | Number nuclear pixels contained between theta -180 < R ≤ -162 | M |
| 255 | Number nuclear pixels contained between theta -162 < R ≤ -144 | M |
| 256 | Number nuclear pixels contained between theta -144 < R ≤ -126 | M |
| 257 | Number nuclear pixels contained between theta -126 < R ≤ -108 | M |
| 258 | Number nuclear pixels contained between theta -108 < R ≤ -90 | M |
| 259 | Number nuclear pixels contained between theta -90 < R ≤ -72 | M |
| 260 | Number nuclear pixels contained between theta -72 < R ≤ -54 | M |
| 261 | Number nuclear pixels contained between theta -54 < R ≤ -36 | M |
| 262 | Number nuclear pixels contained between theta -36 < R ≤ -18 | M |
| 263 | Number nuclear pixels contained between theta -18 < R ≤ 0 | M |
| 264 | Number nuclear pixels contained between theta 0 < R ≤ 18 | M |
| 265 | Number nuclear pixels contained between theta 18 < R ≤ 36 | M |
| 266 | Number nuclear pixels contained between theta 36 < R ≤ 54 | M |
| 267 | Number nuclear pixels contained between theta 54 < R ≤ 72 | M |
| 268 | Number nuclear pixels contained between theta 72 < R ≤ 90 | M |
| 269 | Number nuclear pixels contained between theta 90 < R ≤ 108 | M |
| 270 | Number nuclear pixels contained between theta 108 < R ≤ 126 | M |
| 271 | Number nuclear pixels contained between theta 126 < R ≤ 144 | M |
| 272 | Number nuclear pixels contained between theta 144 < R ≤ 162 | M |
| 273 | Number nuclear pixels contained between theta 162 < R ≤ 180 | M |
| 274 | 0.1 quantile of nuclear pixels from the boundary line | M |
| 275 | 0.2 quantile of nuclear pixels from the boundary line | M |
| 276 | 0.3 quantile of nuclear pixels from the boundary line | M |
| 277 | 0.4 quantile of nuclear pixels from the boundary line | M |
| 278 | 0.5 quantile of nuclear pixels from the boundary line | M |
| 279 | 0.6 quantile of nuclear pixels from the boundary line | M |
| 280 | 0.7 quantile of nuclear pixels from the boundary line | M |

| | | |
|---|---|---|
| 281 | 0.8 quantile of nuclear pixels from the boundary line | M |
| 282 | 0.9 quantile of nuclear pixels from the boundary line | M |
| 283 | 1 quantile of nuclear pixels from the boundary line | M |
| 284 | 0.1 quantile of luminal pixels from the boundary line | M |
| 285 | 0.2 quantile of luminal pixels from the boundary line | M |
| 286 | 0.3 quantile of luminal pixels from the boundary line | M |
| 287 | 0.4 quantile of luminal pixels from the boundary line | M |
| 288 | 0.5 quantile of luminal pixels from the boundary line | M |
| 289 | 0.6 quantile of luminal pixels from the boundary line | M |
| 290 | 0.7 quantile of luminal pixels from the boundary line | M |
| 291 | 0.8 quantile of luminal pixels from the boundary line | M |
| 292 | 0.9 quantile of luminal pixels from the boundary line | M |
| 293 | 1 quantile of luminal pixels from the boundary line | M |
| 294 | 0.1 quantile of mesangial pixels from the boundary line | M |
| 295 | 0.2 quantile of mesangial pixels from the boundary line | M |
| 296 | 0.3 quantile of mesangial pixels from the boundary line | M |
| 297 | 0.4 quantile of mesangial pixels from the boundary line | M |
| 298 | 0.5 quantile of mesangial pixels from the boundary line | M |
| 299 | 0.6 quantile of mesangial pixels from the boundary line | M |
| 300 | 0.7 quantile of mesangial pixels from the boundary line | M |
| 301 | 0.8 quantile of mesangial pixels from the boundary line | M |
| 302 | 0.9 quantile of mesangial pixels from the boundary line | M |
| 303 | 1 quantile of mesangial pixels from the boundary line | M |
| 304 | Luminal textural contrast | T |
| 305 | Luminal textural correlation | T |
| 306 | Luminal textural energy | T |
| 307 | Luminal textural homogeneity | T |
| 308 | PAS+ textural contrast | T |
| 309 | PAS+ textural correlation | T |
| 310 | PAS+ textural energy | T |
| 311 | PAS+ textural homogeneity | T |
| 312 | Nuclear textural contrast | T |
| 313 | Nuclear textural correlation | T |
| 314 | Nuclear textural energy | T |
| 315 | Nuclear textural homogeneity | T |
